# Genome-wide and enzymatic analysis reveals efficient D-galacturonic acid metabolism in the basidiomycete yeast *Rhodosporidium toruloides*

**DOI:** 10.1101/675652

**Authors:** Ryan J. Protzko, Christina A. Hach, Samuel T. Coradetti, Magdalena A. Hackhofer, Sonja Magosch, Nils Thieme, Adam P. Arkin, Jeffrey M. Skerker, John E. Dueber, J. Philipp Benz

**Affiliations:** Department of Molecular and Cell Biology, University of California, Berkeley, California, USA; Energy Biosciences Institute, Berkeley, California, USA; Holzforschung München, TUM School of Life Sciences Weihenstephan, Technische Universität München, Freising, Germany; Environmental Genomics and Systems Biology Division, Lawrence Berkeley National Laboratory, Berkeley, California, USA; Biological Systems & Engineering Division, Lawrence Berkeley National Laboratory, Berkeley, CA 94720, USA; Department of Bioengineering, University of California, Berkeley, California, USA

## Abstract

Biorefining of renewable feedstocks is one of the most promising routes to replace fossil-based products. Since many common fermentation hosts, such as *Saccharomyces cerevisiae*, are naturally unable to convert many component plant cell wall polysaccharides, the identification of organisms with broad catabolism capabilities represents an opportunity to expand the range of substrates used in fermentation biorefinery approaches. The red basidiomycete yeast *Rhodosporidium toruloides* is a promising and robust host for lipid and terpene derived chemicals. Previous studies demonstrated assimilation of a range of substrates, from C5/C6-sugars to aromatic molecules similar to lignin monomers. In the current study, we analyzed *R. toruloides* potential to assimilate D-galacturonic acid, a major sugar in many pectin-rich agricultural waste streams, including sugar beet pulp and citrus peels. D-galacturonic acid is not a preferred substrate for many fungi, but its metabolism was found to be on par with D-glucose and D-xylose in *R. toruloides*. A genome-wide analysis by combined RNAseq/RB-TDNAseq revealed those genes with high relevance for fitness on D-galacturonic acid. While *R. toruloides* was found to utilize the same non-phosphorylative catabolic pathway known from ascomycetes, the maximal velocities of several enzymes exceeded those previously reported. In addition, an efficient downstream glycerol catabolism and a novel transcription factor were found to be important for D-galacturonic acid utilization. These results set the basis for use of *R. toruloides* as a potential host for pectin-rich waste conversions and demonstrate its suitability as a model for metabolic studies in basidiomycetes.

**Importance:** The switch from the traditional fossil-based industry to a green and sustainable bio-economy demands the complete utilization of renewable feedstocks. Many currently used bio-conversion hosts are unable to utilize major components of plant biomass, warranting the identification of microorganisms with broader catabolic capacity and characterization of their unique biochemical pathways. D-galacturonic acid is a plant component of bio-conversion interest and is the major backbone sugar of pectin, a plant cell wall polysaccharide abundant in soft and young plant tissues. The red basidiomycete and oleaginous yeast *Rhodosporidium toruloides* has been previously shown to utilize a range of sugars and aromatic molecules. Using state-of-the-art functional genomic methods, physiological and biochemical assays, we elucidated the molecular basis underlying the efficient metabolism of D-galacturonic acid. This study identifies an efficient pathway for uronic acid conversion to guide future engineering efforts, and represents the first detailed metabolic analysis of pectin metabolism in a basidiomycete fungus.

## Introduction

Negative environmental impacts from fossil fuel consumption and volatile energy costs have accelerated academic and industrial efforts to develop sustainable commodity chemicals and biofuels via microbial fermentation of renewable plant biomass. Pectin-rich side streams from industrial processing of fruits and vegetables have a strong potential as fermentation feedstocks, as they are stably produced in high quantities and can be provided at low cost. Moreover, they accumulate centrally at their respective processing plants, reducing transport costs, are partly pre-treated during the processing, and are naturally devoid of lignin, being a major bottleneck in depolymerization. Furthermore, second generation energy crops, such as agave and sugar beet, have high levels of pectin; sometimes exceeding 40% of the dry weight (1, 2). Despite these major advantages, pectin-rich feedstocks are largely disposed of in landfills and biogas plants or are sold as an inexpensive livestock feed after an energy-intensive drying and pelleting process. Utilizing these waste streams for the biorefinery would benefit the bioeconomy without augmenting current land-use and decrease the contribution of these agricultural wastes to landfill overflow and environmental pollution through airborne spores from molds, which thrive on pectin-rich waste (3, 4).

Pectin is the most heterogeneous of the major plant cell wall polysaccharides and has four main structural classes: homogalacturononan (HG), rhamnogalacturonan I (RG-I), and the substituted HGs rhamnogalacturonan II (RG-II) and xylogalacturonan (XG). α-(1,4)-linked D-galacturonic acid (D-galUA) is the major backbone sugar of all HG structures and can comprise up to 70% of the polysaccharide. D-galUA is a uronic sugar with the same hydroxyl configuration as D-galactose, but with a carboxylic acid group at the C6 position. Other pectic monosaccharides include L-arabinose (L-ara), D-galactose (D-gal), L-rhamnose (L-rha), and D-xylose (D-xyl) (5, 6).

The catabolic pathway for D-galUA utilization has not yet been characterized in the Basidiomycota phylum. In ascomycetes, D-galUA is taken up by an MFS-type transporter specific for uronic acids (7) and in a first step is reduced to L-galactonate by a D-galUA reductase, which is either NADPH-specific or accepts either NADH or NADPH, depending on the organism (8, 9). Next, L-galactonate is transformed into 3-deoxy-L-threo-hex-2-ulosonate by a dehydratase (10) and then into L-glyceraldehyde and pyruvate by an aldolase (11). The last step of the reaction requires NADPH as a cofactor and is catalyzed by a glyceraldehyde reductase which converts L-glyceraldehyde to glycerol, a central metabolite (12).

*Rhodosporidium toruloides* is a strong candidate for bioconversion of pectin-rich waste streams. This basidiomycetous red yeast has been isolated from a wide variety of pectin-rich substrates (e.g. oranges (13), grapes, olives (14) and sugar beet pulp (14, 15). *R. toruloides* can grow well on D-galUA as a sole carbon source (16), indicating an efficient pathway for D-galUA metabolism. Furthermore, *R. toruloides* is of increasing biotechnological interest as a host for bio-conversions. The yeast naturally accumulates lipids and carotenoids, suggesting that it may be a promising host for the production of terpene and lipid-based bio-products (17). Additionally, the yeast can co-utilize both hexose and pentose sugars (18, 19) and assimilate aromatic compounds, such as *p*-coumarate, derived from acylated lignins (20), suggesting advantages for efficient carbon utilization over conventional lignocellulosic conversion hosts. Finally, *R. toruloides* has advantages as a model system for basidiomycetes, as it is easily manipulated in laboratory settings, whereas the basidiomycetes vast majority of known basidiomycetes are difficult to cultivate (21). Furthermore, genetic analyses and mutant strain development are becoming more efficient in *R. toruloides* as novel molecular tools are being developed (22–25).

The aim of the present study was to characterize the D-galUA utilization pathway of *R. toruloides*. Growth assays demonstrate this pathway is highly efficient in comparison to the utilization of D-xyl or even D-glucose (D-glc). To identify all genes involved in D-galUA catabolism, parallel RNAseq and whole-genome RB-TDNAseq studies were performed (22). The enzymes for each metabolic step were subsequently heterologously expressed in *Escherichia coli* and purified to verify their kinetic properties *in vitro*. Furthermore, we identified transporters and a novel transcription factor essential for D-galUA utilization. Finally, global carbon utilization trends underlying the high efficiency of D-galUA catabolism are discussed. We believe the results from this study offer crucial insights into basidiomycete D-galUA utilization and provide a starting point for engineering of *R. toruloides* as a host for pectin-rich waste bioconversion.

## Results

### *R. toruloides* IFO 0880 has a highly efficient D-galUA catabolism and can co-utilize D-galUA with D-glucose and D-xylose

Since it was known that *R. toruloides* can utilize both D-glc and D-xyl (18), we tested how the assimilation of D-galUA would compare to these rates and whether growth inhibition would be visible in mixed substrate cultures. To this end, *R. toruloides* IFO 0880 was grown in 200 μl volume cultures with 50 mM each of these sugars as sole carbon source as well as in cultures in which D-galUA was mixed with either D-glc or D-xyl in a 1:1 ratio. Surprisingly, despite a slightly slower acceleration-phase on D-galUA compared to D-glc in the first 24 hours, culture densities of *R. toruloides* reached almost similar final ODs (Fig. 1A). Moreover, D-galUA was completely consumed by that time, while total consumption of D-glc required about 70 hours (Fig. 1B). With this rate, growth on D-galUA was faster than on D-xyl as the sole carbon source, which required about 80 hours to reach the same density and more than 90 hours to be completely consumed (Fig. 1C,D). In mixed cultures of D-galUA and D-glc, D-glc consumption was accelerated compared to single inoculations, indicating co-utilization of both sugars (Fig. 1B). The same was true for the co-cultures of D-galUA and D-xyl (Fig. 1D). Also in this case, the presence of D-galUA led to an acceleration of D-xyl assimilation, while the D-galUA utilization was slightly delayed compared to the single inoculations.

**Figure 1.**
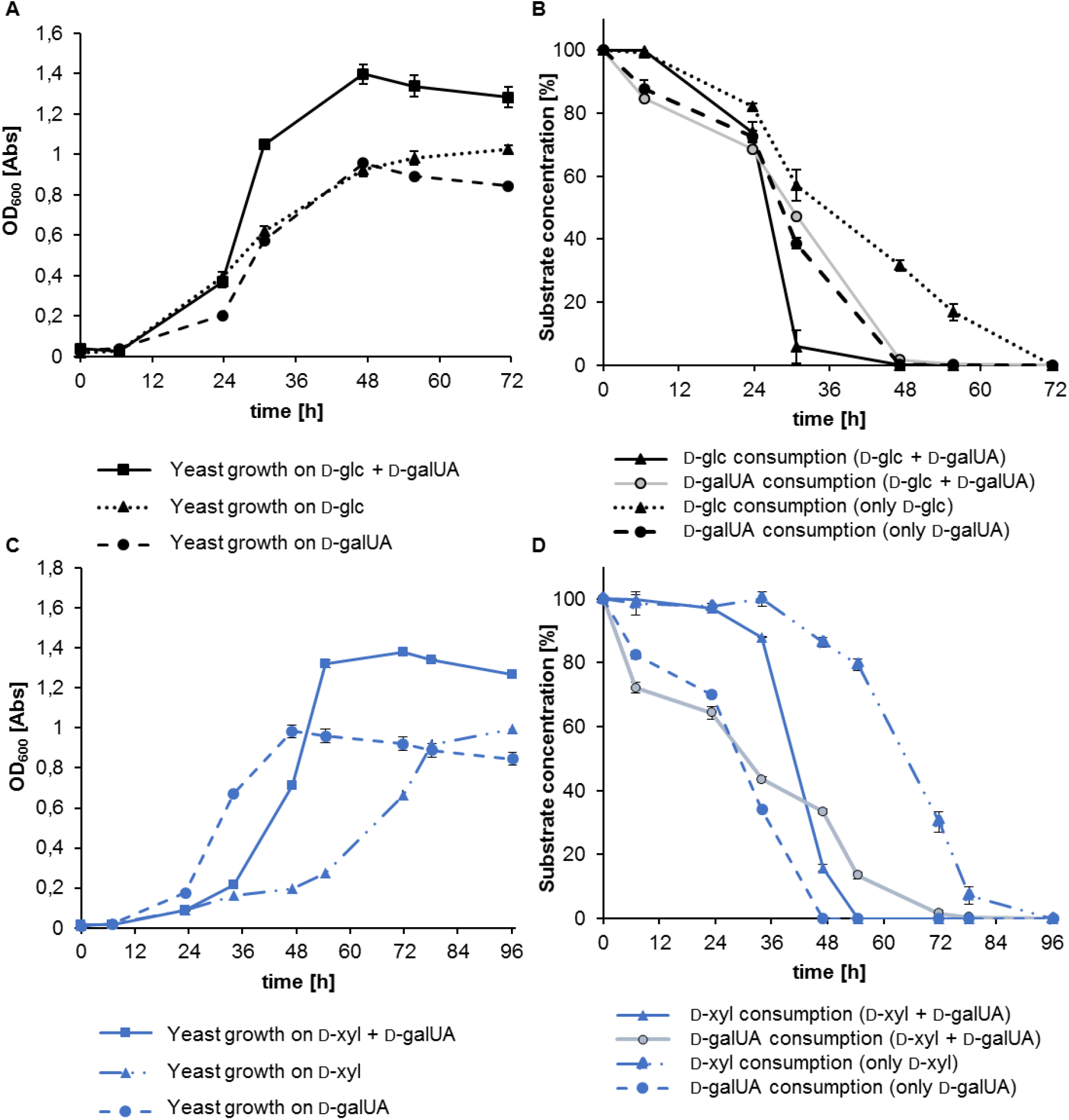
Growth assays of *R. toruloides* IFO 0880 on single carbon source media exhibit efficient utilization of D-galUA compared to D-glu and D-xyl. Growth assays on mixed carbon source media show co-consumption of D-galUA with D-glu and D-galUA with D-xyl. **(A)** Culture growth (OD) over time on 50 mM each of D-galUA, D-glc and 1:1 mixed substrates. **(B)** Normalized sugar consumption of the same cultures. Dotted lines, dashed lines and solid lines indicate D-glc, D-galUA and mixed cultures, respectively. For sugar consumption of mixed cultures, D-glc consumption is shown by a black line, whereas D-galUA consumption is shown in grey. **(C)** Culture growth (OD) over time on 50 mM each of D-galUA, D-xyl and 1:1 mixed substrate. **(D)** Normalized sugar consumption of the same cultures. Dotted lines, dashed lines and solid lines indicate D-xyl, D-galUA and mixed cultures, respectively. For sugar consumption of mixed cultures, D-xyl consumption is shown as a blue line, whereas D-galUA consumption is shown in grey. Values are the mean of three biological replicates. Error bars indicate standard deviation (SD).

Since the above experiments were performed at low concentrations of monosaccharides, we performed an additional growth assay at 500 mM of substrate and larger volume (50 mL) to resemble industrial settings with improved economics (Fig. 2). The high sugar loadings were tolerated by *R. toruloides*, reaching similar ODs like observed in the small cultures for D-galUA and about doubled culture densities on D-glc as the sole carbon source, respectively. The mixed sugar condition led to an accelerated growth rate, corroborating the positive effect of co-consumption that was already visible in the initial assays. These results demonstrate that the presence of D-galUA appears not to be inhibitory to the catabolism of C5 and C6 sugars, but rather leads to enhanced utilization.

**Figure 2.**
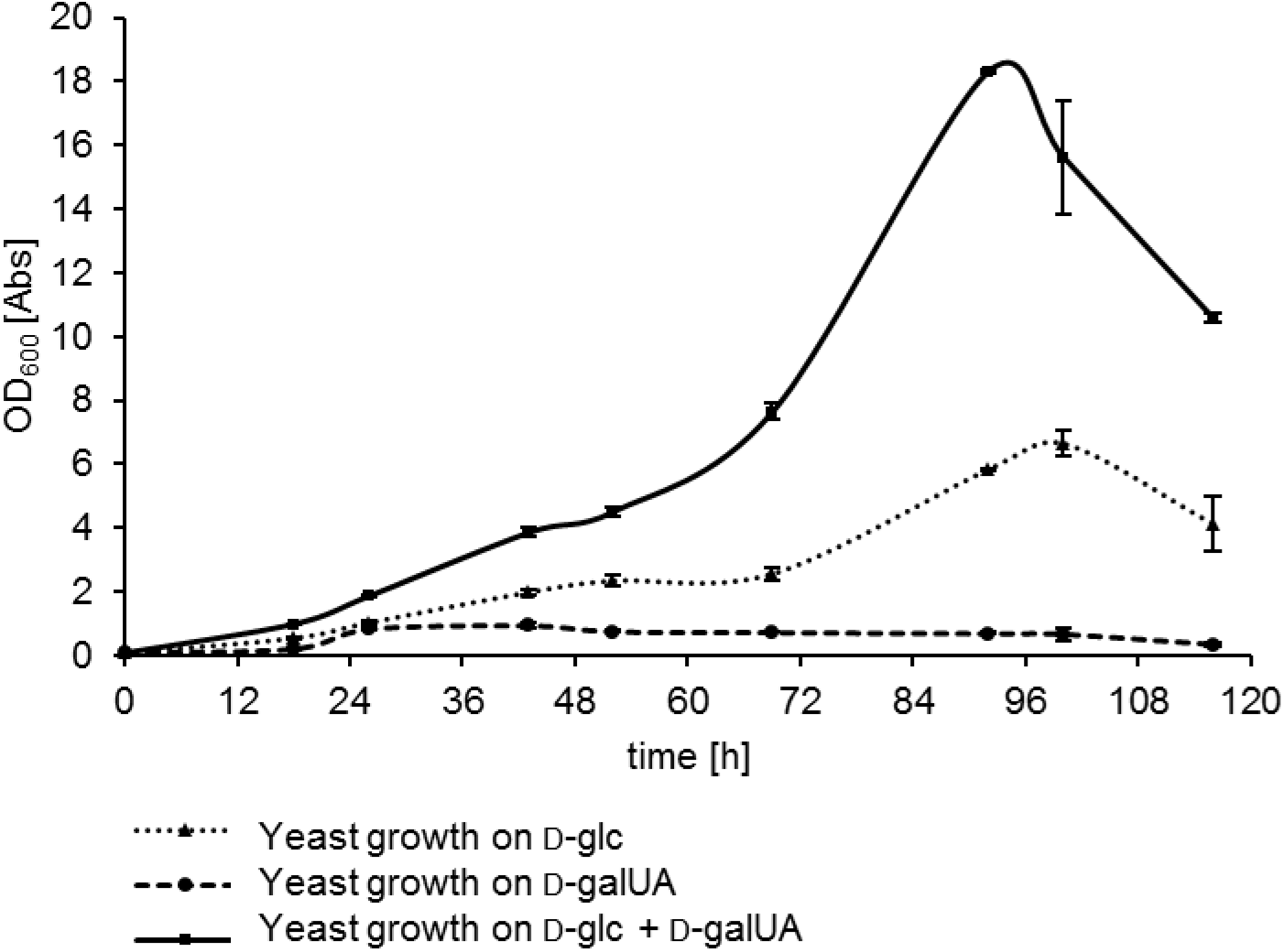
Growth assays of *R. toruloides* IFO 0880 on single and mixed cultures (1:1) of D-glc and D-galUA on 500 mM substrate concentration in 50 mL culture volume. Culture growth (OD) was measured over time. Dotted lines, dashed lines and solid lines indicate D-glc, D-galUA and mixed cultures, respectively. Values are the mean of three biological replicates. Error bars indicate standard deviation (SD).

### Identification of putative D-galUA utilization genes using differential RNAseq analysis

We hypothesized that the genes involved in D-galUA utilization in *R. toruloides* could be identified by analyzing the transcriptional response to media containing D-galUA as the sole carbon source compared to media containing either D-glc or glycerol. Therefore, after growth of *R. toruloides* IFO 0880 on either 2% D-galUA, 2% glycerol or 2% D-glc, RNA was extracted during log growth phase and the transcriptome analyzed by RNAseq. Overall, more than 2,000 genes displayed differential transcript abundances between these three conditions, reflecting the significantly different requirements for growth on these carbon sources (SI Table 1). Growth on D-galUA and glycerol was more alike, in this respect, than growth on D-glc, as can be seen from the condition clustering (Fig. 3). Hierarchical clustering furthermore separated the differentially expressed genes into three clusters of 869 genes most highly expressed on D-glc, 889 genes most highly expressed on glycerol and 625 genes most highly expressed on D-galUA (Fig. 3). This last cluster included several genes with sequence similarity to known enzymes participating in D-galUA catabolism in *Aspergillus niger*, *Trichoderma reesei* and *Neurospora crassa* (Table 1).

**Figure 3.**
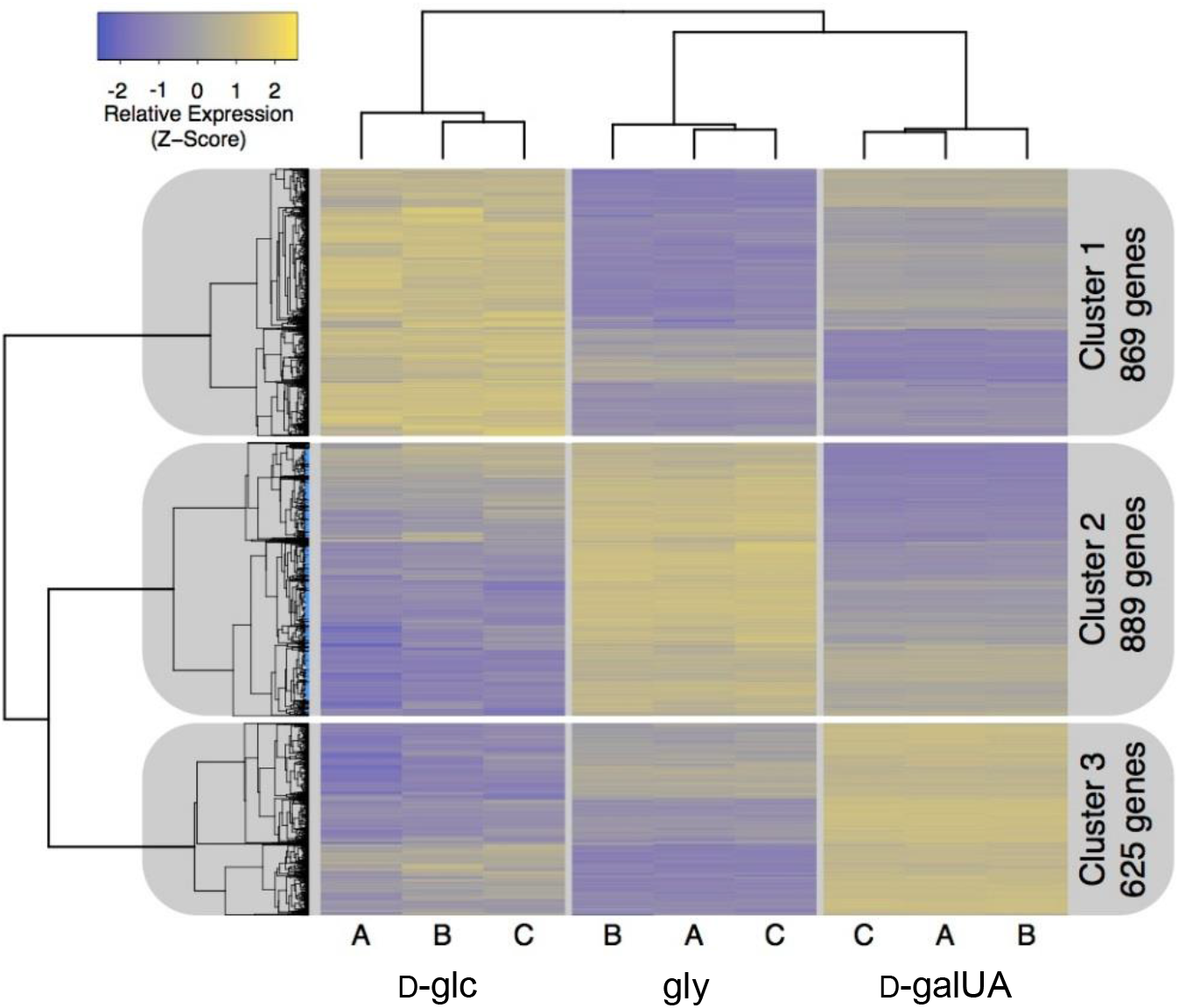
Hierarchical clustering of *R. toruloides* gene expression data reveals three major clusters, based on carbon source. Cluster 1 contains 869 genes most highly expressed on D-glucose (D-glc), Cluster 2, contains 889 genes most highly expressed on glycerol (gly) and Cluster 3 contains 625 genes most highly expressed on D-galacturonate (D-galUA).

**Table 1.** Putative enzymes involved in D-galUA catabolism are upregulated when *R. toruloides* is grown in D-galacturonic acid (D-galUA) as a sole carbon source (Fig. 3, Cluster 3). The RTO4 protein ID, homologs to filamentous fungi D-galUA catabolic pathway genes, reported/putative function, and FPKM (Fragments Per Kilobase of transcript per Million mapped reads) values are shown for D-galUA, D-glucose (D-glc) and glycerol (gly).

| Protein ID | Homolog | Reported/Putative Function | D-galUA | D-glc | gly |
| --- | --- | --- | --- | --- | --- |
| RTO4_12062 | *A. niger* GaaB<br>*T. reesei* Lgd1<br>*N. crassa* NCU07064 | L-galactonic acid dehydratase | 9846 | 26 | 57 |
| RTO4_11882 | *T. reesei* Gar1 | D-galUA reductase | 7667 | 118 | 64 |
| RTO4_12061 | *A. niger* GaaC<br>*T. reesei* Lga1<br>*N. crassa* NCU09532 | 2-keto-3-deoxy-l-galactonate aldolase | 6397 | 24 | 134 |
| RTO4_9774 | *A. niger* GaaD (a.k.a LarA, Alr, Err1)<br>*T. reesei* Gld1 | L-glyceraldehyde reductase<br>pentose reductase, erythrose reductase | 3981 | 524 | 699 |

### Identification of genes required for D-galUA metabolism using genome-wide fitness profiling

To rapidly assess which *R. toruloides* genes are necessary for growth in D-galUA, we grew a sequence-barcoded random insertion library of *R. toruloides* IFO 0880 on either 2% D-galUA, 2% gly or 2% D-glc similar to the RNAseq analysis above. Insertions in genes necessary for growth in the respective carbon sources should prevent or slow growth in those conditions, thus leading to a depletion in the relative abundance of the sequence barcodes associated with those insertions (22). T-DNA insertions in 28 genes led to significant growth defects on D-galUA versus D-glc, and insertions in 20 genes led to significant growth defects on D-galUA versus glycerol (Table 2; Fig. 4). After filtering for statistical significance, we combined our RNAseq data and fitness profiling datasets to gain further insight into the metabolism of D-galUA. Only seven genes had at least a 2-fold increase in transcript abundance on D-galUA and at least a 2-fold decrease in abundance for insertional mutants on D-galUA compared to glycerol and D-glc (Table 2). These genes included homologs to the previously characterized D-galUA utilization pathways in *A. niger* (9) (GaaB, GaaC, and GaaD) and *T. reesei* (GAR1; (8)). RTO4_9841, an MFS-type transporter related to pentose transporters (e.g. LAT-1 in *N. crassa*; NCU02188) was also both transcriptionally induced and required for robust growth on D-galUA, although other transporters were also specifically induced on D-galUA and may therefore be involved in the transport of D-galUA (Table 2). RTO4_13270, a fungal-specific zinc binuclear cluster transcription factor (TF), was also induced by D-galUA, and insertional mutants were severely deficient for growth on D-galUA, suggesting a primary role in regulating expression of D-galUA utilization enzymes. Finally, an ortholog of GAL7 was also induced and required for robust growth on D-galUA.

**Figure 4.**
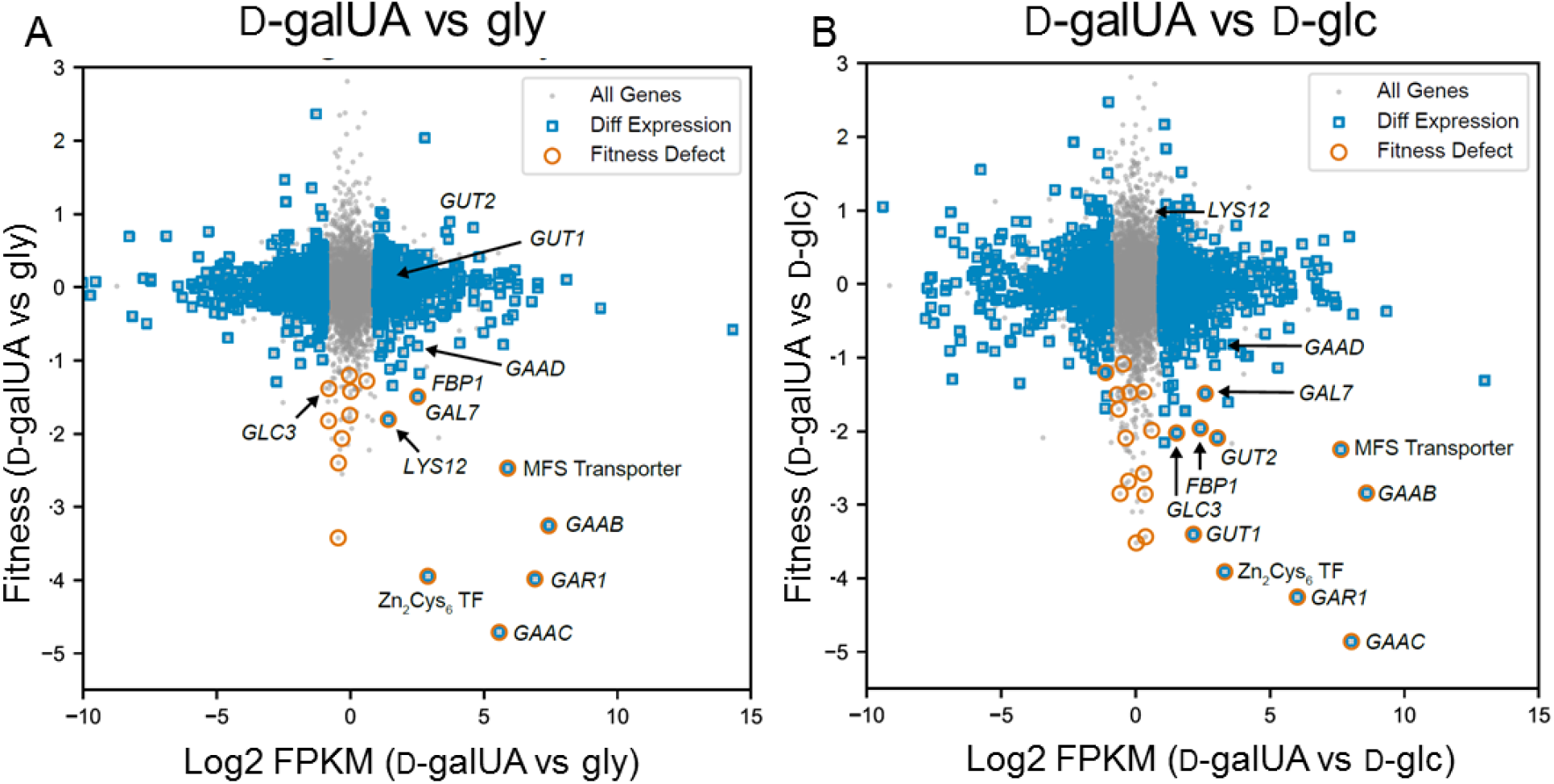
Plotting relative fitness scores *versus* differential expression of D-galUA grown *R. toruloides* identifies genes with essential function in n d-galUA utilization. Genes with significant differential expression had a minimum FPKM > 5, at least a two-fold difference in average FPKM between the two plotted conditions, and a multiple-hypothesis adjusted P-value < 0.05, as calculated across D-galacturonic acid (D-galUA), glycerol (gly), and D-glucose (D-glc) with the Ballgown analysis package for R. Genes with a relative fitness defect had relative T-statistics < −3 between conditions and relative fitness scores < −1 between the two plotted conditions. **(A)** Relative fitness scores vs relative transcript abundance for D-galUA vs gly grown cells. Catabolic pathway genes homologous to the *A. niger* and *T. reesei* utilization pathways (GAR1, GAAB, GAAC and GAAD), a MFS sugar transporter and zinc-finger transcription factor are clearly induced and required for fitness on D-galUA. **(B)** Relative fitness scores vs relative transcript abundance for D-galUA vs D-glc grown cells, illustrating additional pathways involved in D-galUA metabolism. Glycerol catabolism genes, such as GUT1 and GUT2 are found to be induced and required for fitness when grown on D-galUA compared to D-glc.

**Table 2.** Genes with D-galacturonic acid (D-galUA)-specific induction and importance for fitness on D-galUA compared to glycerol (gly) and D-glucose (D-glc) identify a hexose transporter, homologous members to the ascomycete D-galUA catabolism pathway and a putative D-galUA transcription factor. FPKM (Fragments Per Kilobase of transcript per Million mapped reads) and Fitness scores on carbon sources shown.

| Protein ID | Name | Description | Mean FPKM |  |  | Fitness Score |  |  |
| --- | --- | --- | --- | --- | --- | --- | --- | --- |
|  |  |  | D-galUA | gly | D-glc | D-galUA | gly | D-glc |
| RTO4_9841 |  | Hexose transporter | 4605 | 78 | 23 | -2.3 | 0.2 | -0.1 |
| RTO4_11882 | GAR1 | Reductase | 7667 | 64 | 118 | -4.0 | 0.0 | 0.3 |
| RTO4_12062 | GaaB | L-galactonic acid dehydratase | 9846 | 57 | 26 | -3.3 | 0.0 | -0.4 |
| RTO4_12061 | GaaC | 2-keto-3-deoxy-l-galactonate aldolase | 6397 | 134 | 24 | -5.1 | -0.4 | -0.3 |
| RTO4_9774 | GaaD | NADPH-dependent erythrose reductase | 3981 | 699 | 524 | -0.8 | 0.0 | 0.1 |
| RTO4_11332 | GAL7 | Galactose-1-P uridyl transferase | 330 | 58 | 55 | -1.6 | -0.1 | -0.1 |
| RTO4_13270 |  | ZnCys transcription factor | 93 | 12 | 9 | -4.0 | 0.0 | 0.0 |

Additional genes were identified to be required for utilization of D-galUA and glycerol over D-glc (SI Table 2). These genes include members of the canonical glycerol utilization pathway, GUT1 and GUT2, and the glycerol proton symporter, STL1, the latter showing a modest, but statistically significant, growth defect on glycerol. Mutants in homologs to members of the known carbon-catabolite regulating AMPK/SNF1 protein kinase complex (a Mig1/CreA/CRE-1 repressor; SNF1, SNF4, SIP2) (26) were also deficient for growth on one or both of these alternative carbon sources, as were mutants in two G proteins (orthologs of *S. cerevisiae* CDC42 and *H. sapiens* RAB6A) and likely interacting guanine exchange factors. Disruptions in thiolation of some tRNA residues also consistently resulted in small, but significant, fitness defects on D-galUA and glycerol, but not on D-glc, furthering evidence that this process plays a role in nutrient sensing and carbon metabolism in diverse fungi (27, 28).

### *In vitro* enzymatic characterization of the D-galUA catabolic proteins

Based on the data above and previous knowledge from ascomycetes, a model of D-galUA catabolism was hypothesized (Fig. 5). Intriguingly, a two-gene cluster was observed in *R. toruloides*, similar to what was described in ascomycetes (9). However in this case, the gaaC-homolog RTO4_12061 is not linked with the gaaA-homolog (which is absent from the genome), but with RTO4_12062, the homolog of gaaB/lgd1 in *A. niger* and *T. reesei*, respectively (Fig. 5A).

**Figure 5.**
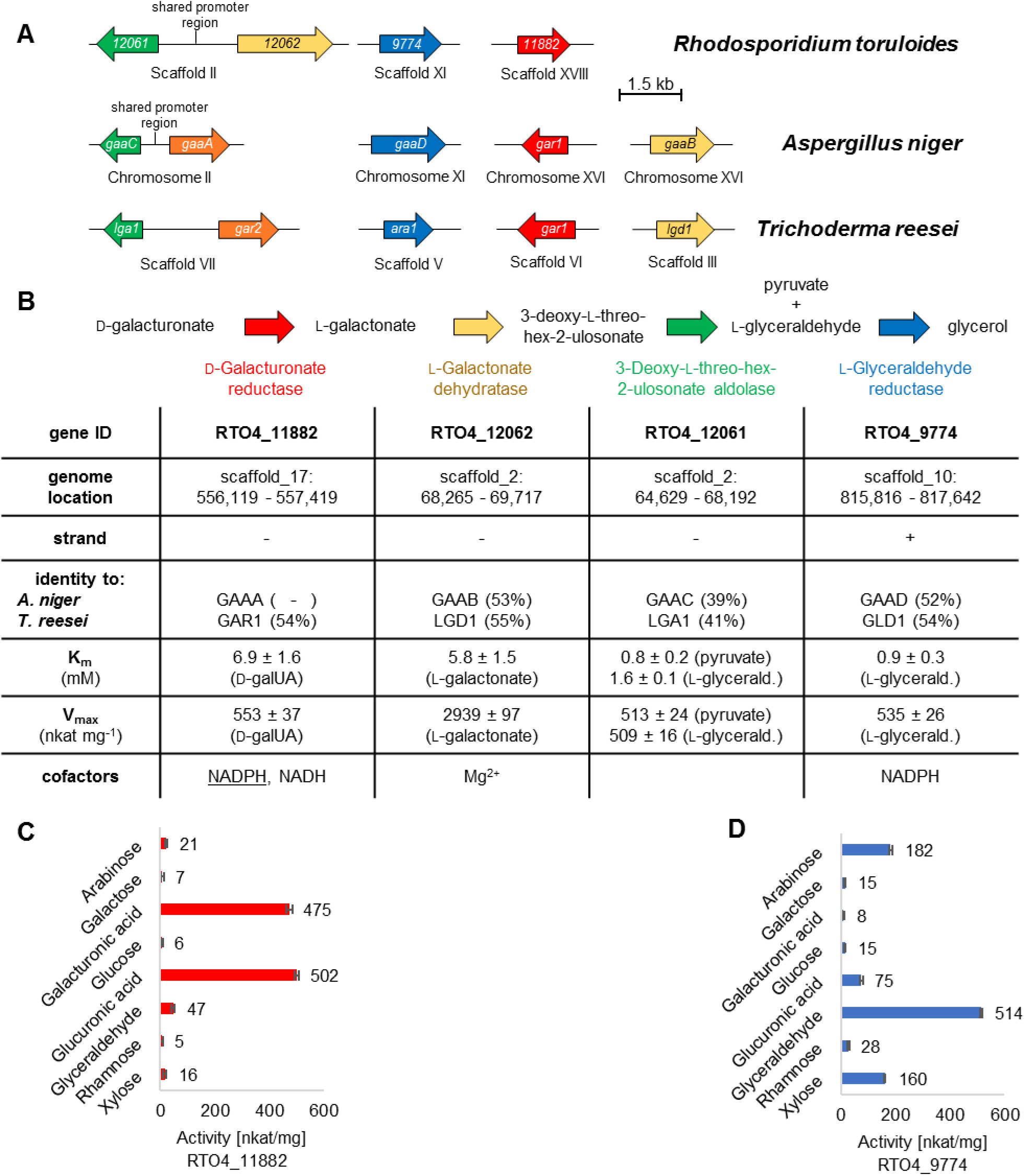
Gene arrangement and kinetic properties of the putative D-galUA metabolizing enzymes in *R. toruloides*. (A) Gene synteny in comparison to *A. niger* and *T. reesei*. In *R. toruloides*, also two genes appear linked in the genome including a shared promoter region, but in contrast to the ascomycetes, it is those coding for the second and third step of the pathway and not the first and third. (B) Table summarizing the major genomic and kinetic properties of the putative D-galUA catabolizing genes and enzymes, respectively. Underlined NADPH indicates the favored cofactor of RTO4_11882. (C) Substrate spectrum of the putative D-galUA reductase. (D) Substrate spectrum of the putative L-glyceraldehyde reductase.

To confirm the corresponding enzyme activities, *in vitro* biochemical studies were performed. The enzymes were heterologously expressed in *E. coli* and purified to characterize their activity. The putative D-galUA reductase and GAR1-homolog RTO4_11882 displayed clear D-galUA reduction activity with a K_M_ of about 7 mM (Fig. 5B; Fig. S1A). The V_max_ at saturating D-galUA concentrations was found to be 553 nkat/mg. A substrate scan revealed similarly high activities for this enzyme also on glucuronic acid with only side activities on all other monosaccharides tested (Fig. 5C), suggesting that RTO4_11882 represents a uronic acid reductase. In addition, the enzyme was found to prefer NADPH as cofactor and showed much weaker activity with NADH (data not shown). Dehydration of L-galactonate by the putative L-galactonate dehydratase (RTO4_12062) was observed at a high V_max_ of 2939 nkat/mg with a K_M_ of 5.8 mM (Fig. 5B; Fig. S1B; (29)). Activity of the putative 3-deoxy-L-threo-hex-2-ulosonate aldolase (RTO4_12061) was tested by monitoring the reverse reaction of L-glyceraldehyde and pyruvate (Fig. 5B; Fig. S1C). Affinities and velocities for both substrates were found to be very similar with a K_M_ in the range of about 1 mM and V_max_ of about 510 nkat/mg (Fig. 5B). The putative L-glyceraldehyde reductase (RTO4_9774) displayed Michaelis Menten kinetics of 0.9 mM with a V_max_ of 535 nkat/mg for L-glyceraldehyde (Fig. 5B; Fig. S1D). A substrate scan interestingly revealed that this enzyme also appears to be a major pentose reductase of *R. toruloides*, since robust activities were found for L-ara and D-xyl with K_M_’s in the range of 20-35 mM (Fig. 5D; Fig. S1D). Lower activities were recorded for other sugars, such as the hexoses D-glc and D-gal, the deoxy-hexose D-rha and the uronic acid D-glucuronic acid.

### Sugar reductase activities are induced by D-galUA in *R. toruloides*

In light of the previous observations, particularly regarding growth physiology and enzymatic characterizations, we aimed to investigate which substrates are able to induce monosaccharide reductase activity in *R. toruloides in vivo*. To this end, reductase assays were performed with whole-cell lysates following growth for 24 h on either D-xyl, D-galUA, D-glc or glycerol. The enzymatic activity of the cell lysates was tested using D-xyl, D-galUA, D-glc and L-glyceraldehyde as substrates by measuring the NADPH concentration loss over time. All lysates showed L-glyceraldehyde reductase activity, albeit to a varying extent (Fig. 6). By contrast, D-galUA and D-glc reductase activity were specific for the cultures grown on D-galUA. Intriguingly, while D-xyl reductase activity was somewhat less specifically induced, growth on D-galUA clearly led to the strongest induction. Considering that RTO4_9774 is induced about 5 to 7-fold on D-galUA over glycerol and D-glc, respectively (Table 1 and 2), these results suggest that a major part of the observed D-xyl reductase activities is contributed by RTO4_9774 functioning as pentose reductase. The same might be true for the low D-glc reductase activity recorded specifically after induction on D-galUA. In addition, these activities could help to explain the accelerated D-xyl and D-glc consumption in presence of D-galUA as seen in the mixed cultures (Fig. 1).

**Figure 6.**
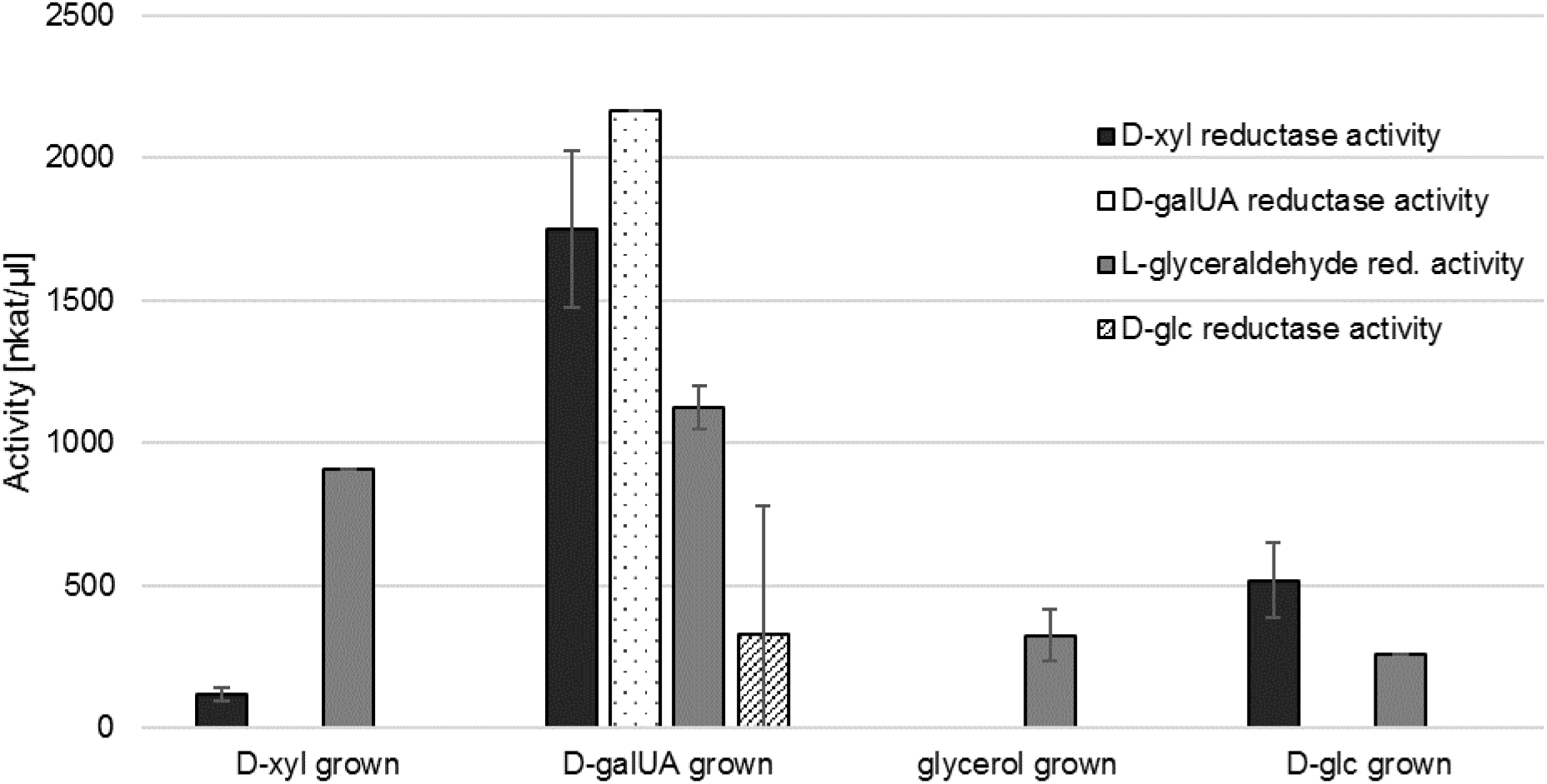
Reductase assays with whole cell lysates of *R. toruloides* IFO 0880 pre-grown on different carbon sources. Measurements were performed from biological triplicate cultures (n=3). Error bars represent standard deviation (SD).

### Identification of RTO4_13270 as the first basidiomycete transcription factor involved in the catabolism of D-galUA

A putative transcription factor, RTO4_13270, exhibited induction on D-galUA (Table 1 and 2). Moreover, disruption of this gene by T-DNA insertion resulted in a very large fitness disadvantage for growth in D-galUA (Table 2 and Fig. 4), indicating that it might be a key regulator for the catabolism of this carbon source. This novel TF belongs to the same Gal4-like family as the known D-galUA-responsive TF from ascomycetes, GaaR (30, 31) but is otherwise phylogenetically unrelated (Fig. 7A). We assessed conservation of RTO4_13270 in basidiomycetes by searching for homologs in the fungal proteomes from the Pucciniomycotina (to which *R. toruloides* belongs; Fig. 7B). Overall, RTO4_13270 was found to be highly conserved in the *Rhodotorula/Rhodosporidium* genus, still relatively conserved in closely related groups, but not at all beyond the Pucciniomycotina.

**Figure 7.**
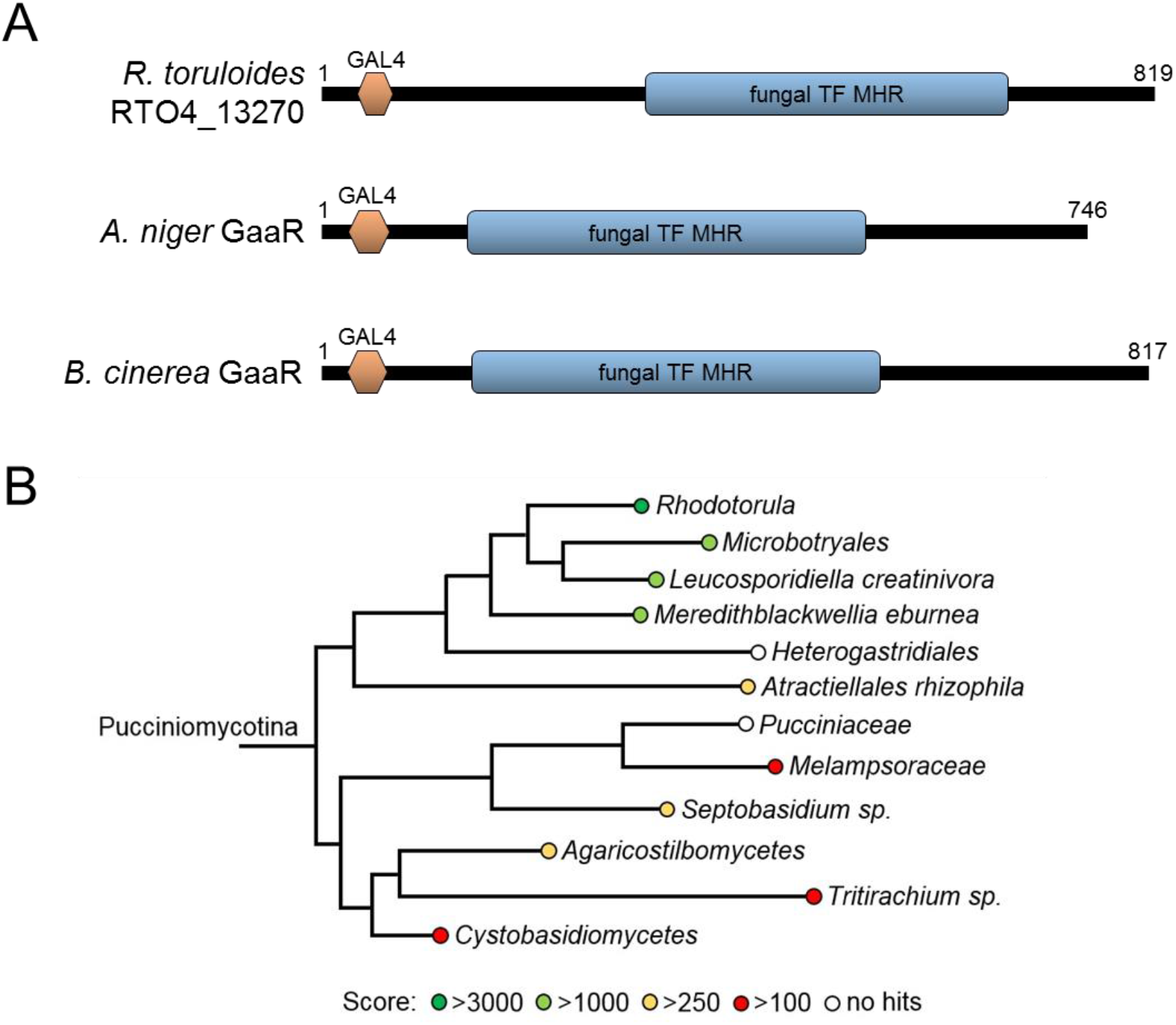
**(A)** Domain analysis of putative D-galUA transcription factors show presence of GAL4 and fungal transcription factor middle homology region (TF MDH) domains; however, RTO4_13270 has low overall homology to the known ascomycete GaaRs. **(B)** Phylogenetic tree of putative RTO4_13270 orthologs. Among the Pucciniomycotina subdivision, proteins most closely related to RTO4_13270 are found in the genus *Leucosporidiella*, *Meredithblackwellia* and *Microbotryales*. To visualize sequence relationships, BLAST Scores were color-coded to the fungal families/genus/species according to their homology. Families with an empty circle did not return any result (these sequences were not publicly available on the Joint Genome Institute’s server).

## Discussion

It has been shown that the red yeast, *R. toruloides*, exhibits strong growth on pectin-derived monosaccharides, including D-galUA, D-xyl, L-ara and D-glc (18). Red yeasts likely fill an opportunistic niche on pectic substrates and assimilate the monosaccharides liberated by the enzymes of other microorganisms (32). For example, *Rhodotorula* species were found to colonize grapes in the presence of other pectinolytic fungi potentially releasing sugars from the fruit tissue (33). In this study, we characterized the D-galUA utilization pathway of *R. toruloides* by a combination of transcriptomics, genome-wide fitness profiling, and biochemical analysis of purified enzymes.

*R. toruloides* utilizes a non-phosphorylative D-galUA catabolic pathway as observed in ascomycete filamentous fungi (Fig. 8) (9). However, the *R. toruloides* pathway is similar to the *T. reesei* pathway compared to the *A. niger* pathway due to the absence of a GaaA homolog and the presence of a functional GAR1 homolog (Fig. 5) (34). The conserved enzymes are highly induced by D-galUA, required for fitness and shown to have the predicted biochemical activities for each catabolic step *in vitro*. When comparing the catalytic activities to those reported for *T. reesei* and *A. niger*, the substrate affinities (K_M_ values) of the *R. toruloides* enzymes are for the most part surprisingly similar (10, 8, 11, 35); Table S3). However, particularly for the dehydratase RTO4_12062, the aldolase RTO4_12061 and the L-glyceraldehyde reductase RTO4_9774, the maximal velocities are about 4 to 500-times higher, suggesting this might contribute to the high efficiency of the pathway flux. Interestingly, the dehydratase and aldolase are clustered in the genome, whereas the pathway reductases are found elsewhere in the genome. Even though this genomic arrangement differs from the state in the filamentous ascomycetes, where the aldolase (*gaaC*) forms a gene pair with the initial D-galUA reductase (*gaaA*) (9), it may indicate that a tight co-regulation of the enzyme expression (via a shared promotor region) is beneficial for the pathway as a whole and for the aldolase in particular.

**Figure 8.**
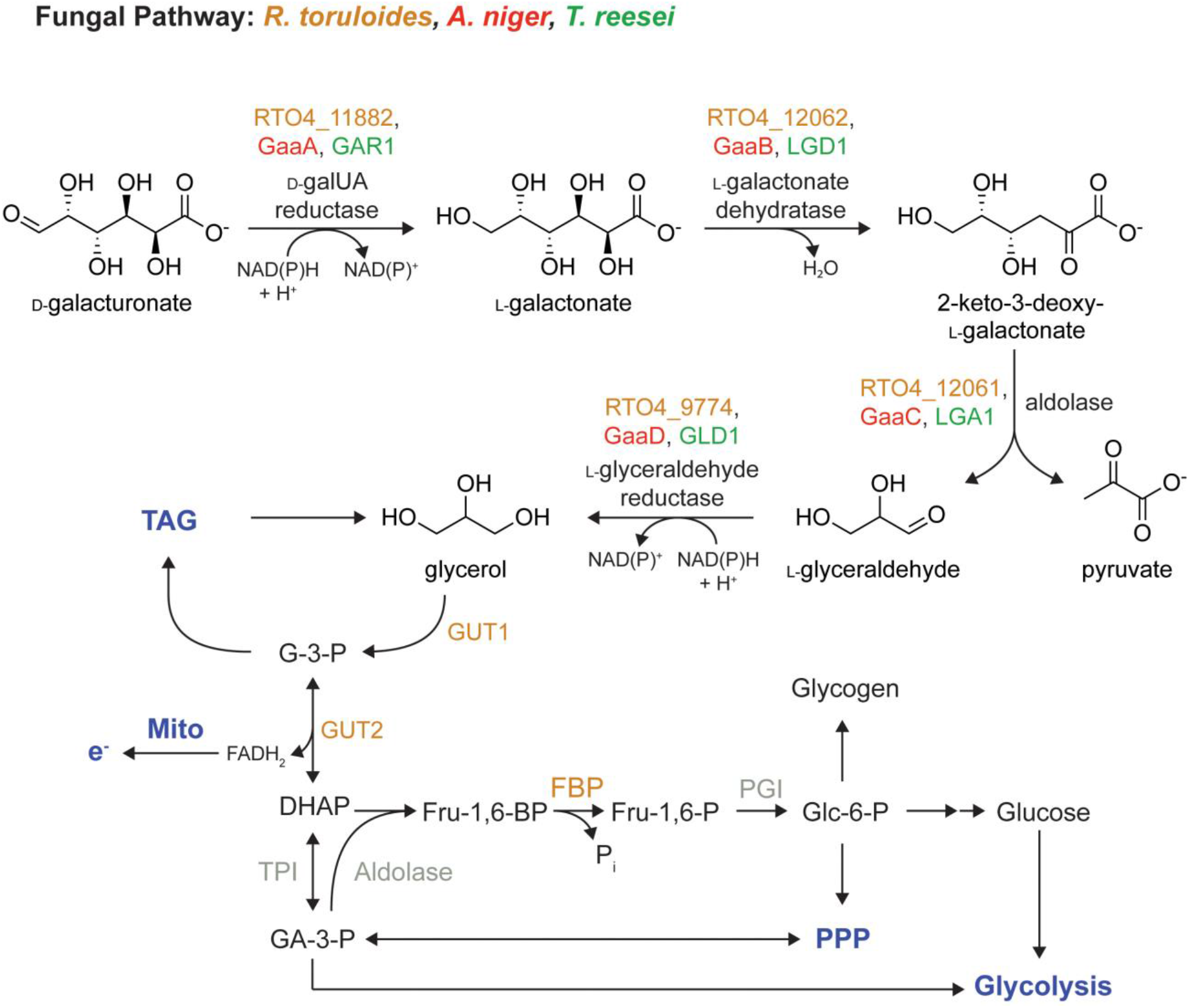
Model of *R. toruloides* D-galUA catabolism based on combined RNAseq and RB-TDNAseq analysis. While the initial enzymatic steps follow the same strategy as known from the ascomycete pathway, genome-wide transcriptional and fitness profiling reveals an expanded role for glycerol metabolism and gluconeogenesis in D-galUA catabolism. TAG (triacylglycerol); Mito (mitochondria); PPP (pentose phosphate pathway).

Another intriguing observation of our study was that the D-galUA catabolism led to an enhanced co-utilization of D-glc and D-xyl, which are the most abundant hexose and pentose sugars in plant biomass and therefore a primary target of biorefinery concepts. This is notable, since it was shown that the presence of D-galUA (at low pH) inhibits the assimilation of D-xyl, D-gal and L-ara in the commonly used fermentation host *S. cerevisiae*, possibly via competitive inhibition of the main transporter Gal2p and a general weak acid toxicity (36). Even though the used pH in this study was higher, the high efficiency catabolism of D-galUA in *R. toruloides* may allow sufficient ATP to overcome intracellular toxicity of pathway intermediates or proton efflux, whereas *cerevisiae* is incapable of D-galUA assimilation without engineering (37–39).

Co-consumption of D-galUA with D-glc and D-xyl might furthermore benefit from the multifunctional role of RTO4_9774 as mentioned above. RTO4_9774 is induced on D-galUA, and its additional activities as a pentose reductase and D-glc reductase will help to assimilate these sugars under mixed culture conditions. Moreover, this competition of sugars for available enzymes may explain the delay in D-galUA consumption. Future experiments with the single gene deletions will help to address this matter. Excitingly, the observation of sugar co-consumption at high sugar loadings is of high biotechnological relevance for efficient mixed-sugar fermentations of pectin-rich biomass.

The contribution of a specific D-galUA uptake system in *R. toruloides* to high flux and low competition with other sugars can be potentially attributed to MFS-type transporters. One MFS-type sugar transporter, RTO4_9841 (class 2.A.1.1; http://www.tcdb.org), exhibited strong induction on D-galUA (mean FPKM of 4605), and disruption of this gene resulted in a significant fitness defect. Interestingly, its close homolog, RTO4_9846, which probably resulted from a recent duplication event, is also induced on D-galUA, but did not cause a significant fitness defect and might therefore represent a pseudogene (see Materials and Methods). Surprisingly, a phylogenetic analysis and comparison with the better described MFS-transporters of class 2.A.1.1 from *N. crassa* (SI Fig. S2) shows that these transporters have higher sequence homology to annotated arabinose transporters (*i.e.* LAT-1; (40), than to the GAT-1 family of D-galUA transporters found in ascomycetes. However, the relation of RTO4_9841 and 9846 to pentose transporters and their function in D-galUA uptake is currently unclear.

The combination of transcriptomics and functional genomics analysis identified downstream components highly relevant for high D-galUA catabolism. Moreover, our emerging model of D-galUA metabolism in *R. toruloides* may also serve as a road map for the engineering of D-galUA pathways in other organisms. In particular, three genes seem to be of high importance and received low fitness scores when disrupted by T-DNA insertions: *GUT1*, *GUT2*, and *FBP1*. The first two genes encode for the enzymes glycerol kinase and mitochondrial glycerol 3-phosphate dehydrogenase that are involved in the canonical glycerol metabolism pathway, as known from *S. cerevisiae* and filamentous fungi (41–43). An efficient glycerol metabolism therefore appears crucial for D-galUA assimilation. Moreover, since *R. toruloides* as an oleaginous yeast has a highly efficient TAG biosynthesis and turnover, which is linked to glycerol metabolism at the stage of glycerol-3-phosphate (G-3-P), the D-galUA metabolism might hitchhike on these capacities and benefit from the high possible fluxes (44, 19). In addition, the FAD-dependent oxidation of G-3-P to dihydroxyacetone phosphate (DHAP) in the mitochondrial outer membrane by GUT2 may provide reducing power necessary for the conversion of relatively oxidized D-galUA. This might be supported by the participation of GUT2 in the G-3-P shuttle, which is involved in the maintenance of the NAD:NADH redox balance for example in *S. cerevisiae* (45). The necessity of the fructose-bisphosphatase (FBP1) for D-galUA utilization may suggest involvement of gluconeogenesis. Further support for this hypothesis might be derived from the essentiality of the SNF1 complex (all three subunits) for growth fitness on D-galUA. This complex is highly conserved from yeast to humans, is an antagonist of carbon catabolite repression, and promotes gluconeogenesis in the absence of D-glc (23). A proper metabolic switch from D-glc to alternative carbon sources such as D-galUA, including activation of gluconeogenesis, is thus a clear prerequisite for efficient growth in this condition. It remains to be shown whether the newly identified TF RTO4_13270 is a target of the SNF1 complex, since one of its main functions is to activate several TFs by phosphorylation in yeast (46, 47). A downstream product of FBP1, glucose-6-phosphate, also represents an entry gate into the pentose phosphate pathway (PPP), which may be an additional way to generate the necessary reducing equivalents to redox-balance the assimilation of D-galUA, an oxidized sugar acid substrate.

The characterization of the D-galUA catabolic pathway described in this work sets the basis for use of *R. toruloides* as potential host for pectin-rich waste conversions. The novel enzymes and transporters described here may furthermore be valuable for biotechnological use in anaerobic microbes, such as *S. cerevisiae*. The present study also demonstrates that the molecular tools now available for *R. toruloides* make it an ideal model for the elucidation of basic biological concepts, such as carbon sensing, signaling and substrate utilization (among many others), in basidiomycete fungi.

## Materials and Methods

### Strains

We used *R. toruloides* IFO0880 (also designated as NBRC 0880), which was obtained from the NITE Biological Resource Center (https://www.nite.go.jp/nbrc/) for the growth assays, transcriptional analysis and functional genomic analysis performed in this study.

### Culture conditions

Unless otherwise stated, *R. toruloides* IFO0880 cultures were grown at 30 °C in 50 mL liquid media in 250 mL baffled flasks with 250 rpm agitation on a shaker. For strain maintenance rich media conditions, yeast peptone dextrose (YPD) media was used supplemented with 2% (w/v) D-glc. For growth assays, IFO0880 was pre-cultured in 0.68% (w/v) yeast nitrogen base w/o amino acids (Sigma-Aldrich, Y0626), pH 5.5 with 2% (w/v) D-glc for 4 days. Afterwards, cells were washed in YNB w/o carbon. YNB cultures supplemented with the respective carbon source and 0.2% (w/v) ammonium sulfate were inoculated with an initial OD_600_ of 0.1. For mixed cultures, substrates were combined in equal amounts to the respective final concentration.

To investigate the growth of *R. toruloides* on different carbon sources, we compared 50 mM D-glc, D-galUA and D-xyl as either a single carbon source or in combination. Furthermore, growth on D-glc and D-galUA was also tested at a higher concentration of 500 mM. Assays performed with 50 mM substrate concentration were performed using sterile 96-well plates with cover. The plates were constantly shaken at 1000 rpm at 30 °C in a thermoblock. For assays with 500 mM substrate concentration, cells were cultured in 50 mL culture volume using 250 mL baffled shake flasks. For transcriptional analysis, *R. toruloides* was grown in media containing either 50 mM D-galUA or glycerol. *E. coli* strains were cultured in LB medium supplemented with the respective antibiotics ampicillin (100 μg/mL) and chloramphenicol (68 μg/mL), incubated at 37 °C and 250 rpm constant agitation.

### Sugar consumption assays

At each time point of absorbance measurement, aliquots were taken and diluted with water to a concentration of 1:100. The samples were centrifuged at for 4 min at 15.000 x g and 4°C. Subsequently, 400 μL of the supernatant was transferred into a new Eppendorf tube. To determine the remaining sugar content, we used High pH Anion Exchange Chromatography (HPAEC). Prior to measuring, aliquots of the supernatants were further diluted to a final concentration of 1:5000. Quantification of monosaccharides was performed as described in (48) and (7). For elution of neutral monosaccharides, samples were injected into a 3 x 150 mm CarboPac PA20 column at 30 °C by using an isocratic mobile phase of 10 mM NaOH and a flowrate of 0.4 mL/min over 13 min.

### RNA sequencing and analysis

An overnight starting culture of *R. toruloides* 0880 was diluted to OD (600 nm) 0.2 in 100 mL YPD (BD Biosciences, BD242820, San Jose, CA) in a 250 mL baffled flask and incubated 8 hours at 30 °C, 200 rpm on a platform shaker. In this time the culture reached an OD 1.0. Cultures were then pelleted by centrifugation at 3000 RCF at room temperature for 5 minutes and washed twice with yeast nitrogen base (YNB) (BD Biosciences, BD291940) media without carbon source. This starter culture was then used to inoculate triplicate cultures in YNB plus 2% D-glc (Sigma-Aldrich, G7528, St. Louis, MO), 2% D-galUA, and 2% glycerol (Sigma-Aldrich, G5516, St. Louis, MO) at OD 0.1 in 100 mL cultures in 250 mL baffled flasks. Each carbon source was then grown to the onset of stationary phase at OD 2.0. Approximately 10 OD units were then pelleted and frozen at −80 °C for DNA extraction and analysis. Total RNA was isolated using the RNeasy Mini Kit (Qiagen, Cat No. 74104) using on column DNA digestion (Qiagen, Cat No. 79254). RNAseq libraries were sequenced on an Illumina HiSeq 4000 system at the QB3 Vincent J. Coates Genomic Sequencing Laboratory (http://qb3.berkeley.edu/gsl/) using standard mRNA enrichment, library construction and sequencing protocols. Approximately 40 million 50 bp single-end reads were acquired per replicate per condition. Transcript abundances and differential expression were calculated with HiSat 2.1.0, StringTie 1.3.3b, and Ballgown 2.8.4 (49) by mapping against *R. toruloides* IFO 0880 v4 reference transcripts (https://genome.jgi.doe.gov/Rhoto_IFO0880_4/Rhoto_IFO0880_4.home.html).

To cluster genes with significant differential expression, genes with adjusted P-values < 0.05 across all three conditions were filtered to remove genes with low expression (FPKM < 5 in all conditions) and small fold changes (< 2-fold). FPKM values were then clustered using Pearson correlation as the similarity metric and average linkage as the clustering method (hclust function R)

### Barcode sequencing and Fitness Analysis

Fitness analysis of a pooled, barcoded *R. toruloides* T-DNA mutant library was performed as described in (22) with minor alterations. Briefly, three aliquots of the previously generated pool of random *Agrobacterium tumefaciens* insertional mutants were thawed on ice then inoculated at OD (600 nm) 0.2 in 100 mL YPD (BD Biosciences, BD242820, San Jose, CA) in a 250 mL baffled flask and incubated 8 hours at 30 °C, 200 rpm on a platform shaker. In this time the mutant pools reached an OD 0.8. At this time a ‘time 0’ reference sample of the starter culture was collected for all three replicates. Cultures were then pelleted by centrifugation at 3000 RCF at room temperature for 5 minutes and washed twice with yeast nitrogen base (YNB) (BD Biosciences, BD291940) media without carbon source. Each starter culture was then used to inoculate new cultures in YNB plus 2% D-glc (Sigma-Aldrich, G7528, St. Louis, MO), 0.2% D-glc, 2% D-galUA, 0.2% D-galUA, and 2% glycerol (Sigma-Aldrich, G5516, St. Louis, MO) at OD 0.1 in 100 mL cultures in 250 mL baffled flasks. Each carbon source was then grown to the onset of stationary phase (14, 14, 36, 19, and 64 hours respectively, with OD at sampling of 5, 1.3, 4.1, 0.8 and 1.9 respectively). Approximately 10 OD units were then pelleted and frozen at −80 °C for DNA extraction and analysis. Barcode amplification, sequencing and fitness analysis were performed as described in (22).

### Lysate assay

Cells were pre-cultured for 2 days in 3 mL YNB (pH 5.5) plus 2 % D-glc in 24-well deep well plates at 700 rpm agitation and 30 °C. Cells were washed 3 times in 1 mL YNB medium without carbon source, followed by 24 h of induction in 3 mL YNB with the respective carbon source. A volume of 1 mL of the culture was centrifuged at 3.500 rpm at 4 °C for 2 min to lyse the cells. The cell pellet was disrupted in 400 μL of lysis buffer (50 mM NaOH, 1 mM EDTA, 1% Triton X-100) with 0.5 mm glass beads using a laboratory bead mill (BeadBug Microtube homogenizer, SLG Gauting) for 1 min at maximum speed and RT. For the determination of reductase activity by measuring the loss of NADPH concentration over time, the complete supernatant after cell lysis was used. The total reaction mixture of 100 μL contained 50 mM substrate (D-xyl, D-glc, D-galUA or glycerol), 0.3 mM NADPH, 100 mM sodium phosphate buffer pH 7, 0.1 μL of Tween20 and 50 μL of the different cell lysates. The assay was performed at 30 °C in UV-compatible 96-well plates (Corning, Germany) in an Infinite® M200 PRO reader (Tecan, Germany) for 5 min in total, measuring the optical density at 340 nm every 15 sec. Prior to each measurement, the plates were shaken for 5 sec.

### *In vitro* activity assays

For activity determination of *in vitro* purified enzymes, RTO4_11882, RTO4_12062, RTO4_12061, and RTO4_9774 were cloned into a custom-made HIS6-TEV expression vector under control of a T7 promoter and transformed into *E. coli* (Rosetta) cells. Protein expression was induced by using 1 mM IPTG at OD_600_ 0.4 - 0.6 and cells incubated overnight at 16 °C and 250 rpm agitation before harvesting by centrifugation at 4 °C. For lysis, cells were resuspended in lysis buffer (see above), 20 mg/mL lysozyme was added and the suspension incubated for 1 h at 37 °C rotating at 10 rpm. After 30 min and 45 min of incubation, 2.5 μL of DNase I was added. Cell debris was precipitated by centrifugation at 16.000 x g and 4 °C for 20 min. The supernatant was used for protein purification by immobilized metal-affinity chromatography (IMAC) (50). Chosen elutions were desalted in Vivaspin columns (10,000 Da MW cutoff; Sartorius, Germany) with storage buffer (20 mM sodium phosphate, 20 mM NaCl, pH 7) by centrifugation at 3.000 x g and 4 °C.

The affinity tag of the putative L-galactonate dehydratase was removed by incubation with an in-house purified and His-tagged TEV protease (ratio 40:1 (mg:mg)) overnight at 4 °C. In a second IMAC, the tag-free protein was collected in the flow through and desalted as described above. Protein concentrations were determined with a Bradford assay using Roti®-Quant reagent (Roth, Germany) and a BSA calibration series. The enzymatic activity of the two reductases was determined as described above, but with a total reaction volume of 200 μl and 1 – 2 μg purified protein instead of cell lysates.

For RTO4_12061, a thiobarbiturate assay according to (51) was performed. To this end, 200 μl reaction mix (50 mM sodium phosphate buffer pH 7.0, 60 mM pyruvate and varying L-glyceraldehyde concentrations or 25 mM L-glyceraldehyde and varying pyruvate concentrations) were mixed with 3.3 μg protein. Samples were removed during the linear range of the reaction, stopped and developed as described, but with half the quantities.

For RTO4_12062, a modified semicarbazide assay was performed (29). 1 μg enzyme was mixed with different L-galactonate concentrations in 5 mM MgCl_2_ and 50 mM sodium phosphate (pH 7.0) in 180 μl total volume. Within the linear range of the reaction, 80 μl samples were taken, mixed with 20 μl 2 M HCl, vortexed, and centrifuged at 16.000 x g and 4 °C for 10 min. 40 μl of the supernatant were pipetted to 160 μl semicarbazide solution (1 % (w/v) semicarbazide, 1 % (w/v) sodium acetate) and incubated at RT for 30 min. The absorbance was measured at 250 nm in UV-compatible 96-well plates (Corning, Germany). Linear L-galactonate was obtained by dissolving L-galactono-1,4-lactone (Sigma, Germany) in water, and adding sodium hydroxide until the pH stopped changing and could be set to pH 7 with 1 M sodium phosphate.

### Synteny identification

Protein sequences of GAR1, GAR2, LGD1, LGA1 and GLD1 from *Trichoderma reesei* were acquired from The Universal Protein Resource (UniProt; https://www.uniprot.org/) databases. Sequences were compared by BLASTp (https://blast.ncbi.nlm.nih.gov/Blast.cgi) with *T. reesei* sequences from FungiDB (http://fungidb.org/fungidb/) and the results illustrating 100% sequence identity were chosen for determination of scaffolds, strand orientation (+ or - strand) and genomic DNA sequence (from ATG to STOP codon). *Aspergillus niger* protein sequences of gaaA, gaaB, gaaC, gaaD were acquired from FungiDB. The GAR2 sequence of *reesei* (from Uniprot) was used to determine the *A. niger* ortholog in FungiDB by using BLASTp. Their strand orientation (+ or – strand), position on the chromosomes, as well as their genomic DNA sequence (from ATG to STOP codon) was determined in the FungiDB. For *R. toruloides* IFO_0880, protein sequences of RTO4_11882, RTO4_12062, RTO4_12061 and RTO4_9774 were acquired from MycoCosm (http://jgi.doe.gov/fungi) using the *R. toruloides* IFO 0880 v4 genome. Their strand orientation (+ or – strand), position on the scaffolds, as well as their genomic DNA sequence (from ATG to STOP codon) was determined in MycoCosm. In general, the DNA sequence length of all genes was determined to create an approximation of the gene length in the respective figure. The amino acid sequence of transcription factor RTO4_13270 was retrieved from JGI database (https://genome.jgi.doe.gov/Rhoto_IFO0880_4/Rhoto_IFO0880_4.home.html) and compared to all entries in the Pucciniomycotina tree of MycoCosm by using BLASTp. The best hit of each sub-group was taken and their BLAST score was used to determine homology. The phylogeny of the Pucciniomycotina was taken from MycoCosm and modified with the results from the homology search.

### Phylogenetic analyses

The protein sequences of RTO4_11882, RTO4_12062, RTO4_12061 and RTO4_9774 from IFO 0880 v4 were retrieved from MycoCosm (http://jgi.doe.gov/fungi) and compared to their homologs in *T. reesei* and *A. niger*. Additionally, the conservation of these genes between *R. toruloides* and representative basidiomycete organisms was determined by phylogenetic analysis. Amino acid sequences of *T. reesei* and *A. niger* were retrieved from UniProt and FungiDB, respectively, as described in the previous section. The protein sequences of the basidiomycete organisms were retrieved from MycoCosm by running BLASTp of the *R. toruloides* genes against the Basidiomycota tree. Genes with high sequence similarity and from representative families were used for the following analyses. The phylogenetic tree was constructed using Clustal Omega (https://www.ebi.ac.uk/Tools/msa/clustalo/) (52–54). The design was finalized using FigTree v1.4.3 (http://tree.bio.ed.ac.uk/software/figtree/) to create a rectangular tree with increasing node ordering and branches transformed to cladogram.

Similarly, *R. toruloides* proteins annotated as sugar porters or MFS-type by Pfam (http://pfam.xfam.org/) (55–59) were analyzed using the transporter categories database (TCDB; http://www.tcdb.org/) (60). Amino acid sequences of *R. toruloides* proteins classified as category 2.A.1.1 (sugar porter family) were retrieved from MycoCosm and compared to sequences of *N. crassa* OR74a transporter proteins of TCDB category 2.A.1.1 (60), which were gathered from FungiDB. For RTO4_9846, due to doubts about correct annotation, an alternative gene model was proposed and used for the phylogenetic analysis. The gene model can be found at: https://genome.jgi.doe.gov/cgi-bin/dispGeneModel?db=Rhoto_IFO0880_4&id=9846. As above, a phylogenetic tree was created with Clustal Omega using these sequence data. The tree was finalized using FigTree v1.4.3 with a polar tree layout, increasing node ordering and branches transformed to cladogram.

## Acknowledgements

We thank Petra Arnold, Nicole Ganske, Nadine Griesbacher and Sabrina Paulus (HFM, TUM) for excellent technical assistance. We furthermore thank Jonas Andrich and Luke N. Latimer for help to start the project (UCB).

The following funding is gratefully acknowledged: R.J.P. was supported by an NSF graduate fellowship. J.E.D. was supported by NSF Award 1330914. J.M.S and A.P.A (UCB) were supported by Energy Biosciences Institute grants OO1605 and OO6J01 as well as US Dept. of Energy (Office of Biological and Environmental Research) grant DE-SC-0012527.

---

**SI Table S2.**
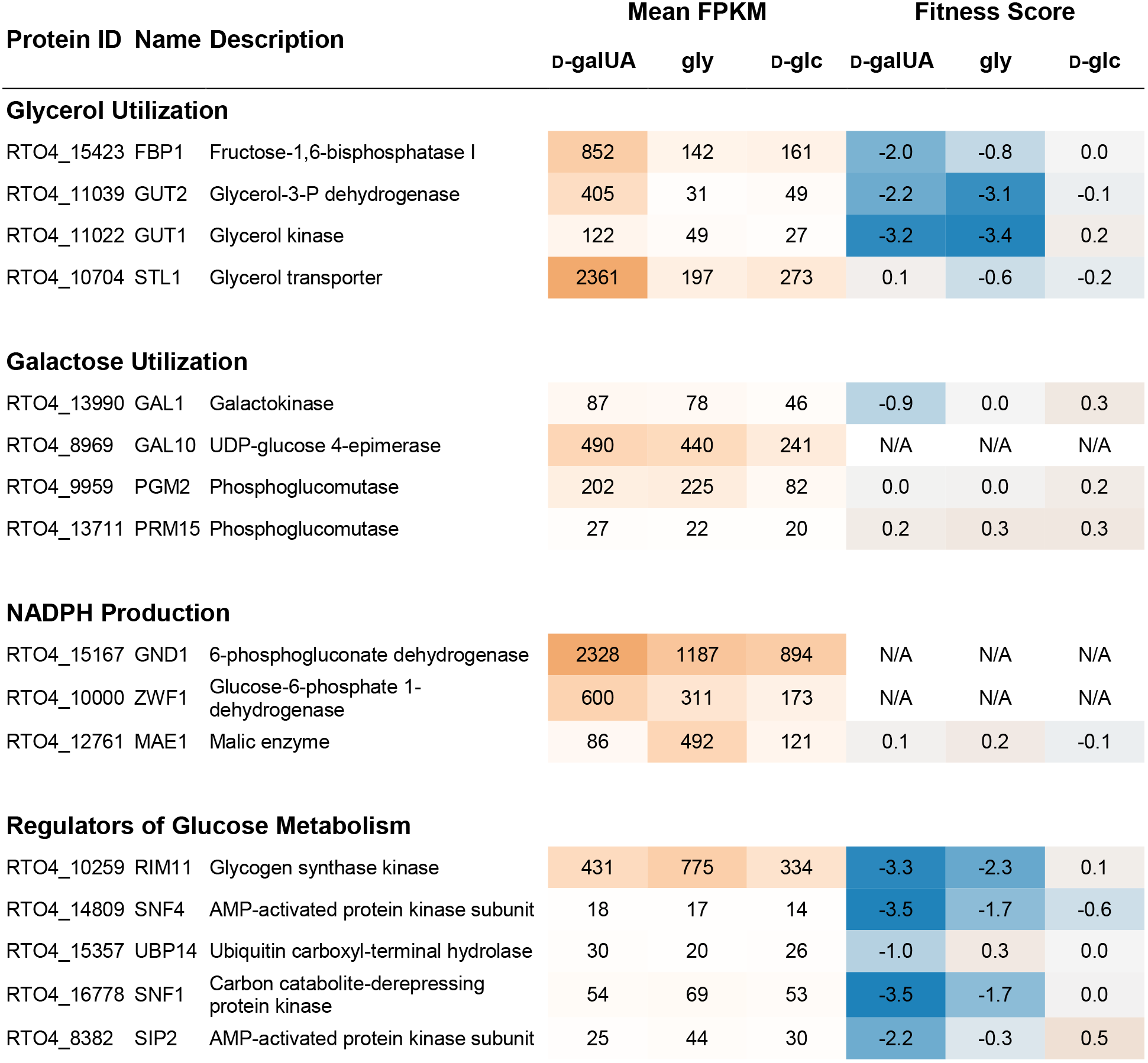
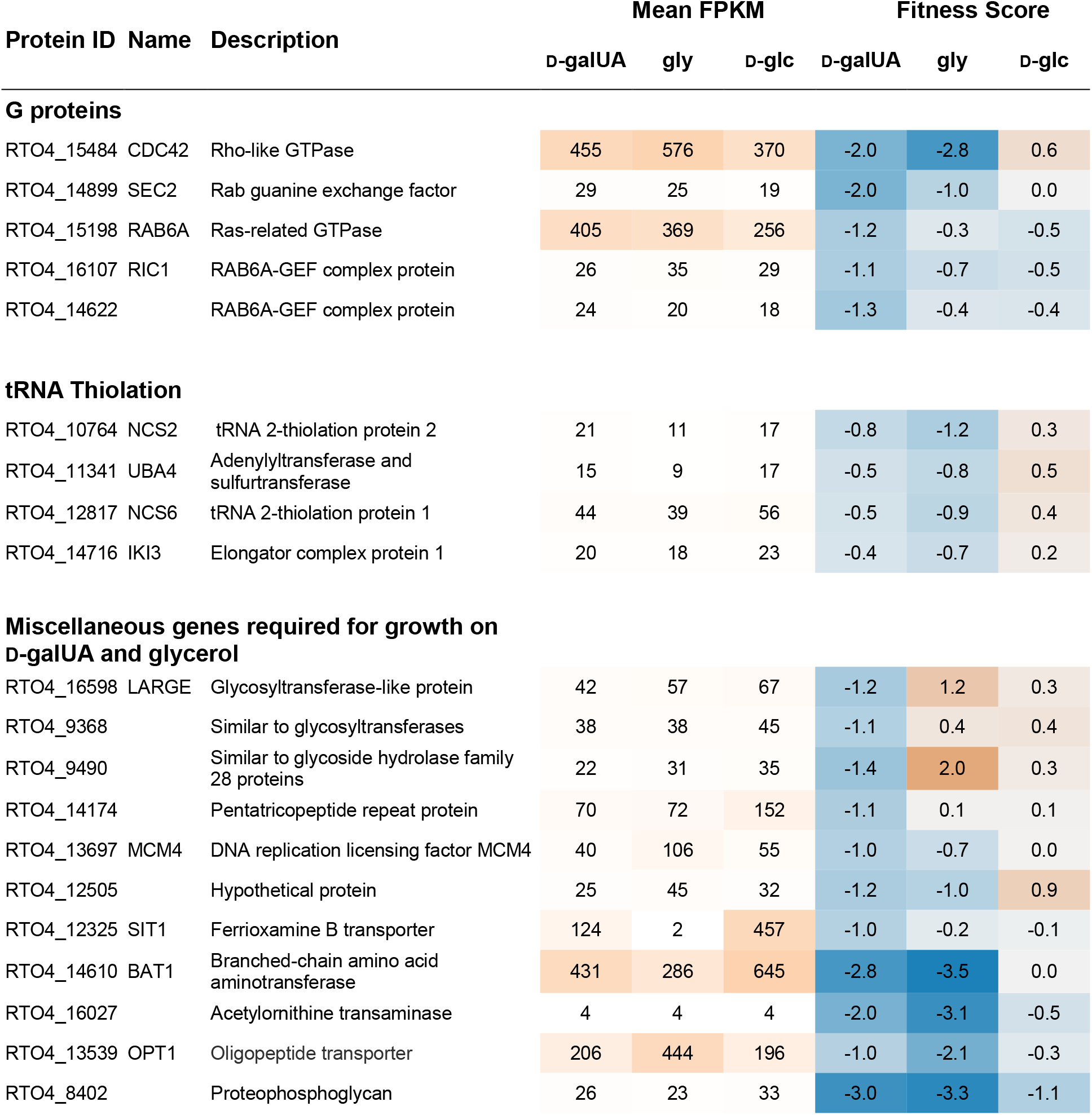
Genes from potentially interesting functional groups with fitness defects and/or induction on D-galacturonic acid (D-galUA) and glycerol (gly) compared to D-glucose (D-glc) identifies global genetic factors required for respective carbon source utilization. Essential genes do not have fitness scores (N/A) as they are absent from the insertion library. FPKM values are shaded orange in proportion expression level. Fitness scores shaded blue in proportion to negative scores and orange for positive scores. Genes with significant fitness defects and high induction on D-galacturonic acid (D-galUA) and glycerol (gly) compared to D-glucose (D-glc) identifies global genetic factors required for respective carbon source utilization. Essential genes do not have fitness scores (N/A) as they are absent from the insertion library. FPKM values are shaded orange in proportion expression level. Fitness scores shaded blue in proportion to negative scores and orange for positive scores.

| Protein ID | Name | Description | Mean FPKM |  |  | Fitness Score |  |  |
| --- | --- | --- | --- | --- | --- | --- | --- | --- |
|  |  |  | D-galUA | gly | D-glc | D-galUA | gly | D-glc |
| Glycerol Utilization |  |  |  |  |  |  |  |  |
| RTO4_15423 | FBP1 | Fructose-1,6-bisphosphatase I | 852 | 142 | 161 | -2.0 | -0.8 | 0.0 |
| RTO4_11039 | GUT2 | Glycerol-3-P dehydrogenase | 405 | 31 | 49 | -2.2 | -3.1 | -0.1 |
| RTO4_11022 | GUT1 | Glycerol kinase | 122 | 49 | 27 | -3.2 | -3.4 | 0.2 |
| RTO4_10704 | STL1 | Glycerol transporter | 2361 | 197 | 273 | 0.1 | -0.6 | -0.2 |
| Galactose Utilization |  |  |  |  |  |  |  |  |
| RTO4_13990 | GAL1 | Galactokinase | 87 | 78 | 46 | -0.9 | 0.0 | 0.3 |
| RTO4_8969 | GAL10 | UDP-glucose 4-epimerase | 490 | 440 | 241 | N/A | N/A | N/A |
| RTO4_9959 | PGM2 | Phosphoglucomutase | 202 | 225 | 82 | 0.0 | 0.0 | 0.2 |
| RTO4_13711 | PRM15 | Phosphoglucomutase | 27 | 22 | 20 | 0.2 | 0.3 | 0.3 |
| NADPH Production |  |  |  |  |  |  |  |  |
| RTO4_15167 | GND1 | 6-phosphogluconate dehydrogenase | 2328 | 1187 | 894 | N/A | N/A | N/A |
| RTO4_10000 | ZWF1 | Glucose-6-phosphate 1-dehydrogenase | 600 | 311 | 173 | N/A | N/A | N/A |
| RTO4_12761 | MAE1 | Malic enzyme | 86 | 492 | 121 | 0.1 | 0.2 | -0.1 |
| Regulators of Glucose Metabolism |  |  |  |  |  |  |  |  |
| RTO4_10259 | RIM11 | Glycogen synthase kinase | 431 | 775 | 334 | -3.3 | -2.3 | 0.1 |
| RTO4_14809 | SNF4 | AMP-activated protein kinase subunit | 18 | 17 | 14 | -3.5 | -1.7 | -0.6 |
| RTO4_15357 | UBP14 | Ubiquitin carboxyl-terminal hydrolase | 30 | 20 | 26 | -1.0 | 0.3 | 0.0 |
| RTO4_16778 | SNF1 | Carbon catabolite-derepressing protein kinase | 54 | 69 | 53 | -3.5 | -1.7 | 0.0 |
| RTO4_8382 | SIP2 | AMP-activated protein kinase subunit | 25 | 44 | 30 | -2.2 | -0.3 | 0.5 |
|  |  |  | D-galUA | gly | D-glc | D-galUA | gly | D-glc |
| G proteins |  |  |  |  |  |  |  |  |
| RTO4_15484 | CDC42 | Rho-like GTPase | 455 | 576 | 370 | -2.0 | -2.8 | 0.6 |
| RTO4_14899 | SEC2 | Rab guanine exchange factor | 29 | 25 | 19 | -2.0 | -1.0 | 0.0 |
| RTO4_15198 | RAB6A | Ras-related GTPase | 405 | 369 | 256 | -1.2 | -0.3 | -0.5 |
| RTO4_16107 | RIC1 | RAB6A-GEF complex protein | 26 | 35 | 29 | -1.1 | -0.7 | -0.5 |
| RTO4_14622 |  | RAB6A-GEF complex protein | 24 | 20 | 18 | -1.3 | -0.4 | -0.4 |
| tRNA Thiolation |  |  |  |  |  |  |  |  |
| RTO4_10764 | NCS2 | tRNA 2-thiolation protein 2 | 21 | 11 | 17 | -0.8 | -1.2 | 0.3 |
| RTO4_11341 | UBA4 | Adenylyltransferase and sulfurtransferase | 15 | 9 | 17 | -0.5 | -0.8 | 0.5 |
| RTO4_12817 | NCS6 | tRNA 2-thiolation protein 1 | 44 | 39 | 56 | -0.5 | -0.9 | 0.4 |
| RTO4_14716 | IKI3 | Elongator complex protein 1 | 20 | 18 | 23 | -0.4 | -0.7 | 0.2 |
| Miscellaneous genes required for growth on D-galUA and glycerol |  |  |  |  |  |  |  |  |
| RTO4_16598 | LARGE | Glycosyltransferase-like protein | 42 | 57 | 67 | -1.2 | 1.2 | 0.3 |
| RTO4_9368 |  | Similar to glycosyltransferases | 38 | 38 | 45 | -1.1 | 0.4 | 0.4 |
| RTO4_9490 |  | Similar to glycoside hydrolase family 28 proteins | 22 | 31 | 35 | -1.4 | 2.0 | 0.3 |
| RTO4_14174 |  | Pentatricopeptide repeat protein | 70 | 72 | 152 | -1.1 | 0.1 | 0.1 |
| RTO4_13697 | MCM4 | DNA replication licensing factor MCM4 | 40 | 106 | 55 | -1.0 | -0.7 | 0.0 |
| RTO4_12505 |  | Hypothetical protein | 25 | 45 | 32 | -1.2 | -1.0 | 0.9 |
| RTO4_12325 | SIT1 | Ferrioxamine B transporter | 124 | 2 | 457 | -1.0 | -0.2 | -0.1 |
| RTO4_14610 | BAT1 | Branched-chain amino acid aminotransferase | 431 | 286 | 645 | -2.8 | -3.5 | 0.0 |
| RTO4_16027 |  | Acetylornithine transaminase | 4 | 4 | 4 | -2.0 | -3.1 | -0.5 |
| RTO4_13539 | OPT1 | Oligopeptide transporter | 206 | 444 | 196 | -1.0 | -2.1 | -0.3 |
| RTO4_8402 |  | Proteophosphoglycan | 26 | 23 | 33 | -3.0 | -3.3 | -1.1 |

**SI Table S3.** Comparison of catalytic activities of conserved enzymes in the non-phosphorylative D-galUA catabolic pathway between *R. toruloides*, *T. reesei* and *A. niger*.

| Protein | Substrate(s) | Activity<br>*R. toruloides* | Activity<br>*T. reesei* | Activity<br>*A. niger* | Ref. |
| --- | --- | --- | --- | --- | --- |
| D-Galacturonate reductase | D-galUA;<br>NADPH<br>$K_M$<br>$V_{Max}$ | **RTO4_11882**<br>6.9 ± 1.6 mM<br>553 ± 37 nkat/mg | **GAR1**<br>6 mM<br>666.8 nkat/mg | **GaaA**<br>0.175 mM<br>145 nkat/mg | (9, 8, this study) |
| L-Galactonate dehydratase | L-galactonate<br>$K_M$<br>$V_{Max}$ | **RTO4_12062**<br>5.8 mM<br>2939 ± 97 nkat/mg | **LGD1**<br>5 mM<br>- (cell lysate) | **GaaB**<br>3.4 mM<br>5.6 nkat/mg | (9, 8, this study) |
| 3-Deoxy-L-threo-hex-2-ulosonate aldolase | pyruvate<br>$K_M$<br>$V_{Max}$ | **RTO4_12061**<br>0.8 ± 0.2 mM<br>513 ± 24 nkat/mg | **LGA1**<br>0.5 mM<br>108.4 nkat/mg | **GaaC**<br>-<br>- | (11, this study) |
| | L-glycerald.<br>$K_M$<br>$V_{Max}$ | **RTO4_12061**<br>1.6 ± 0.1 mM<br>509 ± 16 nkat/mg | **LGA1**<br>1.2 mM<br>108.4 nkat/mg | **GaaC**<br>-<br>- | (11, this study) |
| L-Glycer-aldehyde reductase | L-glycerald.<br>$K_M$<br>$V_{Max}$ | **RTO4_9774**<br>0.9 ± 0.3 mM<br>535 ± 26 nkat/mg | **GLD1**<br>0.9 mM<br>140 nkat/mg | **GaaD**<br>-<br>- | (35, this study) |
| | L-arabinose<br>$K_M$<br>$V_{Max}$ | **RTO4_9774**<br>20.1 mM ± 5.0 mM<br>279 ± 14 nkat/mg | **GLD1**<br>-<br>- | **LarA**<br>54 ± 6 mM<br>500.1 nkat/mg | (61, this study) |
| | D-xylose<br>$K_M$<br>$V_{Max}$ | **RTO4_9774**<br>35.3 ± 5.7 mM<br>294 ± 13 nkat/mg | **GLD1**<br>334 mM<br>450 nkat/mg | **LarA**<br>155 ± 15 mM<br>516.8 nkat/mg | (35, 61, this study) |

**SI Figure S1.**
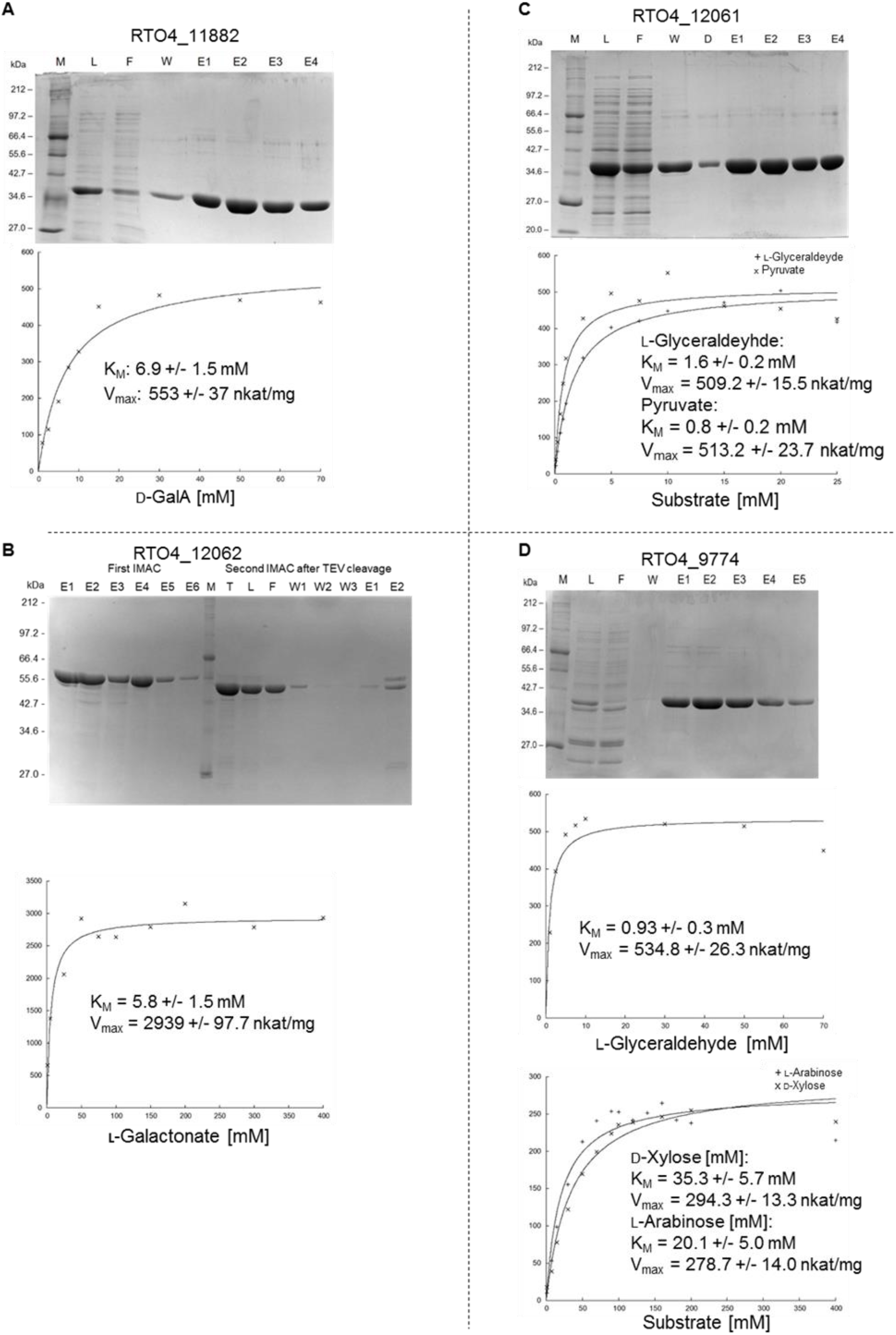
Protein purification and Michaelis Menten kinetics of the recombinant D-galUA catabolism pathway enzymes. His-tagged proteins were expressed in *E. coli* and purified via IMAC; expected molecular weight and purity were determined by SDS-PAGE. Lane descriptions: M = 2 - 212 kDa broad range protein ladder (NEB, Germany), L = load, F = flow, W = wash, T = TEV digest, E = elutions. *In vitro* enzymatic assays were performed to determine the Michaelis Menten kinetics of the enzymes. (A) The activity of RTO4_11882 with an expected protein size of 37.1 kDa was assessed by measuring the loss of NADPH over time. (B) Recombinant RTO4_12062 has an expected protein size of 55.7 kDa (his-tagged) or 53.1 kDa (untagged after TEV cleavage). Activity was determined by a semicarbazide assay. (C) RTO4_12061 has a predicted size of 36.4 kDa and the *in vitro* activity was determined in the reverse direction with the thiobarbiturate (TBA) assay. The two substrates were L-glyceraldehyde and pyruvate. (D) For RTO4_9774, with an expected protein size of 38.0 kDa, the NADPH loss during reduction of L-glyceraldehyde, L-arabinose and D-xylose was measured. Shown data points are the mean of triplicate measurements (n=3). Kinetics data were plotted with the IC50 Tool Kit (http://www.ic50.tk/).

**SI Figure S2:**
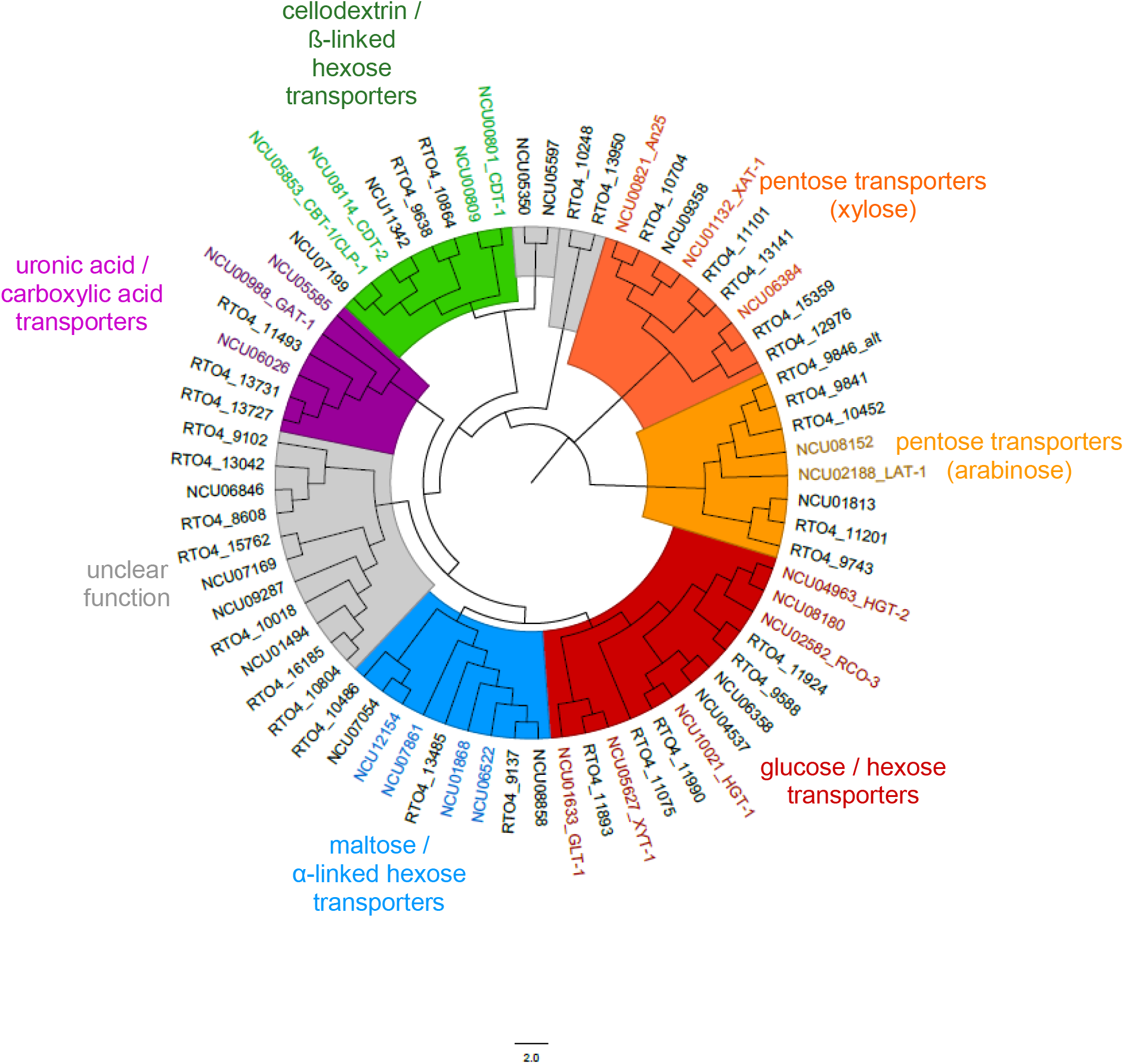
MFS transporter phylogeny using the predicted MFS-type transporters from TCDB-class 2.A.1.1 (sugar porter family) and those from the reference organism *Neurospora crassa* to help in the identification of putative substrates.

## References

1. Linskens HF, Jackson JF. 1999. Analysis of Plant Waste Materials. Modern Methods of Plant Analysis, vol. 20. Springer Berlin Heidelberg, Berlin, Heidelberg.

2. Vendruscolo F, Albuquerque PM, Streit F, Esposito E, Ninow JL. 2008. Apple pomace: a versatile substrate for biotechnological applications. Crit Rev Biotechnol 28:1–12. doi:10.1080/07388550801913840.

3. Schlosser O, Huyard A, Cartnick K, Yañez A, Catalán V, Quang ZD. 2009. Bioaerosol in composting facilities: occupational health risk assessment. Water Environ Res 81:866–877.

4. Wéry N. 2014. Bioaerosols from composting facilities-a review. Front Cell Infect Microbiol 4:42. doi:10.3389/fcimb.2014.00042.

5. Mohnen D. 2008. Pectin structure and biosynthesis. Curr Opin Plant Biol 11:266–277. doi:10.1016/j.pbi.2008.03.006.

6. Atmodjo MA, Hao Z, Mohnen D. 2013. Evolving views of pectin biosynthesis. Annu Rev Plant Biol 64:747–779. doi:10.1146/annurev-arplant-042811-105534.

7. Benz JP, Protzko RJ, Andrich JM, Bauer S, Dueber JE, Somerville CR. 2014. Identification and characterization of a galacturonic acid transporter from *Neurospora crassa* and its application for *Saccharomyces cerevisiae* fermentation processes. Biotechnol Biofuels 7:20. doi:10.1186/1754-6834-7-20.

8. Kuorelahti S, Kalkkinen N, Penttilä M, Londesborough J, Richard P. 2005. Identification in the mold *Hypocrea jecorina* of the first fungal D-galacturonic acid reductase. Biochemistry 44:11234–11240. doi:10.1021/bi050792f.

9. Martens-Uzunova ES, Schaap PJ. 2008. An evolutionary conserved D-galacturonic acid metabolic pathway operates across filamentous fungi capable of pectin degradation. Fungal Genet Biol 45:1449–1457. doi:10.1016/j.fgb.2008.08.002.

10. Kuorelahti S, Jouhten P, Maaheimo H, Penttilä M, Richard P. 2006. L-galactonate dehydratase is part of the fungal path for D-galacturonic acid catabolism. Mol Microbiol 61:1060–1068. doi:10.1111/j.1365-2958.2006.05294.x.

11. Hilditch S, Berghäll S, Kalkkinen N, Penttilä M, Richard P. 2007. The missing link in the fungal D-galacturonate pathway: identification of the L-threo-3-deoxy-hexulosonate aldolase. J Biol Chem 282:26195–26201. doi:10.1074/jbc.M704401200.

12. Sealy-Lewis HM, Fairhurst V. 1992. An NADP+-dependent glycerol dehydrogenase in *Aspergillus nidulans* is inducible by D-galacturonate. Current Genetics 22:293–296. doi:10.1007/BF00317924.

13. Zheng XD, Zhang HY, Sun P. 2005. Biological control of postharvest green mold decay of oranges by *Rhodotorula glutinis*. Eur Food Res Technol 220:353–357. doi:10.1007/s00217-004-1056-5.

14. Vaughn R, Jakubczyk T, MacMillan J, Higgins TE, Dave B, Crampton VM. 1969. Some pink yeasts associated with softening of olives. Appl Microbiol:771–775.

15. Aksu Z, Eren AT. 2005. Carotenoids production by the yeast *Rhodotorula mucilaginosa*: Use of agricultural wastes as a carbon source. Process Biochem 40:2985–2991. doi:10.1016/j.procbio.2005.01.011.

16. Sitepu I, Selby T, Lin T, Zhu S, Boundy-Mills K. 2014. Carbon source utilization and inhibitor tolerance of 45 oleaginous yeast species. J Ind Microbiol Biotechnol 41:1061–1070. doi:10.1007/s10295-014-1447-y.

17. Zhuang X, Kilian O, Monroe E, Ito M, Tran-Gymfi MB, Liu F, Davis RW, Mirsiaghi M, Sundstrom E, Pray T, Skerker JM, George A, Gladden JM. 2019. Monoterpene production by the carotenogenic yeast *Rhodosporidium toruloides*. Microb Cell Fact 18:54. doi:10.1186/s12934-019-1099-8.

18. Wiebe MG, Koivuranta K, Penttilä M, Ruohonen L. 2012. Lipid production in batch and fed-batch cultures of *Rhodosporidium toruloides* from 5 and 6 carbon carbohydrates. BMC Biotechnol 12:26. doi:10.1186/1472-6750-12-26.

19. Bommareddy RR, Sabra W, Maheshwari G, Zeng A-P. 2015. Metabolic network analysis and experimental study of lipid production in *Rhodosporidium toruloides* grown on single and mixed substrates. Microb Cell Fact 14:36. doi:10.1186/s12934-015-0217-5.

20. Yaegashi J, Kirby J, Ito M, Sun J, Dutta T, Mirsiaghi M, Sundstrom ER, Rodriguez A, Baidoo E, Tanjore D, Pray T, Sale K, Singh S, Keasling JD, Simmons BA, Singer SW, Magnuson JK, Arkin AP, Skerker JM, Gladden JM. 2017. *Rhodosporidium toruloides*: a new platform organism for conversion of lignocellulose into terpene biofuels and bioproducts. Biotechnol Biofuels 10:241. doi:10.1186/s13068-017-0927-5.

21. Hansson G, Seifert G. 1987. Effects of cultivation techniques and media on yields and morphology of the basidiomycete *Armillaria mellea*. Appl Microbiol Biotechnol 26. doi:10.1007/BF00253534.

22. Coradetti ST, Pinel D, Geiselman GM, Ito M, Mondo SJ, Reilly MC, Cheng Y-F, Bauer S, Grigoriev IV, Gladden JM, Simmons BA, Brem RB, Arkin AP, Skerker JM. 2018. Functional genomics of lipid metabolism in the oleaginous yeast *Rhodosporidium toruloides*. Elife 7. doi:10.7554/eLife.32110.

23. Park Y-K, Nicaud J-M, Ledesma-Amaro R. 2018. The engineering potential of *Rhodosporidium toruloides* as a workhorse for biotechnological applications. Trends Biotechnol 36:304–317. doi:10.1016/j.tibtech.2017.10.013.

24. Liu X, Zhang Y, Liu H, Jiao X, Zhang Q, Zhang S, Zhao ZK. 2019. RNA interference in the oleaginous yeast *Rhodosporidium toruloides*. FEMS Yeast Res. doi:10.1093/femsyr/foz031.

25. Otoupal PB, Ito M, Arkin AP, Magnuson JK, Gladden JM, Skerker JM. 2019. Multiplexed CRISPR-Cas9-Based Genome Editing of *Rhodosporidium toruloides*. mSphere 4. doi:10.1128/mSphere.00099-19.

26. Sanz P, Viana R, Garcia-Gimeno MA. 2016. AMPK in Yeast: The SNF1 (Sucrose Non-fermenting 1) Protein Kinase Complex. Exp Suppl 107:353–374. doi:10.1007/978-3-319-43589-3_14.

27. Damon JR, Pincus D, Ploegh HL. 2015. tRNA thiolation links translation to stress responses in *Saccharomyces cerevisiae*. Mol Biol Cell 26:270–282. doi:10.1091/mbc.E14-06-1145.

28. Laxman S, Sutter BM, Wu X, Kumar S, Guo X, Trudgian DC, Mirzaei H, Tu BP. 2013. Sulfur amino acids regulate translational capacity and metabolic homeostasis through modulation of tRNA thiolation. Cell 154:416–429. doi:10.1016/j.cell.2013.06.043.

29. Kim S, Lee SB. 2005. Identification and characterization of *Sulfolobus solfataricus* D-gluconate dehydratase: a key enzyme in the non-phosphorylated Entner– Doudoroff pathway. Biochem J 387:271–280. doi:10.1042/BJ20041053.

30. Zhang L, Lubbers RJM, Simon A, Stassen JHM, Vargas Ribera PR, Viaud M, van Kan JAL. 2016. A novel Zn2 Cys6 transcription factor BcGaaR regulates D-galacturonic acid utilization in *Botrytis cinerea*. Mol Microbiol 100:247–262. doi:10.1111/mmi.13314.

31. Alazi E, Niu J, Kowalczyk JE, Peng M, Aguilar Pontes MV, van Kan JAL, Visser J, Vries RP de, Ram AFJ. 2016. The transcriptional activator GaaR of *Aspergillus niger* is required for release and utilization of D-galacturonic acid from pectin. FEBS Lett 590:1804–1815. doi:10.1002/1873-3468.12211.

32. Hackhofer M. 2017. Molecular and biochemical characterization of the pectionolytic capabilities of two basidiomycete red yeasts: *Rhodotorula mucilagionosa* and *Rhodosporidium toruloides*. Master’s thesis.

33. Fleet GH. 2003. Yeast interactions and wine flavour. Int J Food Microbiol 86:11–22.

34. Richard P, Hilditch S. 2009. D-galacturonic acid catabolism in microorganisms and its biotechnological relevance. Appl Microbiol Biotechnol 82:597–604. doi:10.1007/s00253-009-1870-6.

35. Liepins J, Kuorelahti S, Penttilä M, Richard P. 2006. Enzymes for the NADPH-dependent reduction of dihydroxyacetone and D-glyceraldehyde and L-glyceraldehyde in the mould *Hypocrea jecorina*. FEBS J 273:4229–4235. doi:10.1111/j.1742-4658.2006.05423.x.

36. Huisjes EH, Hulster E de, van Dam JC, Pronk JT, van Maris AJA. 2012. Galacturonic acid inhibits the growth of *Saccharomyces cerevisiae* on galactose, xylose, and arabinose. Appl Environ Microbiol 78:5052–5059. doi:10.1128/AEM.07617-11.

37. van Maris AJA, Abbott DA, Bellissimi E, van den Brink J, Kuyper M, Luttik MAH, Wisselink HW, Scheffers WA, van Dijken JP, Pronk JT. 2006. Alcoholic fermentation of carbon sources in biomass hydrolysates by *Saccharomyces cerevisiae*: current status. Antonie Van Leeuwenhoek 90:391–418. doi:10.1007/s10482-006-9085-7.

38. Edwards MC, Doran-Peterson J. 2012. Pectin-rich biomass as feedstock for fuel ethanol production. Appl Microbiol Biotechnol 95:565–575. doi:10.1007/s00253-012-4173-2.

39. Protzko RJ, Latimer LN, Martinho Z, Reus E de, Seibert T, Benz JP, Dueber JE. 2018. Engineering *Saccharomyces cerevisiae* for co-utilization of D-galacturonic acid and D-glucose from citrus peel waste. Nat Commun 9:5059. doi:10.1038/s41467-018-07589-w.

40. Benz JP, Chau BH, Zheng D, Bauer S, Glass NL, Somerville CR. 2014. A comparative systems analysis of polysaccharide-elicited responses in *Neurospora crassa* reveals carbon source-specific cellular adaptations. Mol Microbiol 91:275–299. doi:10.1111/mmi.12459.

41. Hondmann DH, Busink R, Witteveen CF, Visser J. 1991. Glycerol catabolism in *Aspergillus nidulans*. J Gen Microbiol 137:629–636. doi:10.1099/00221287-137-3-629.

42. Klein M, Swinnen S, Thevelein JM, Nevoigt E. 2017. Glycerol metabolism and transport in yeast and fungi: established knowledge and ambiguities. Environ Microbiol 19:878–893. doi:10.1111/1462-2920.13617.

43. Ho P-W, Swinnen S, Duitama J, Nevoigt E. 2017. The sole introduction of two single-point mutations establishes glycerol utilization in *Saccharomyces cerevisiae* CEN.PK derivatives. Biotechnol Biofuels 10:10. doi:10.1186/s13068-016-0696-6.

44. Zhu Z, Zhang S, Liu H, Shen H, Lin X, Yang F, Zhou YJ, Jin G, Ye M, Zou H, Zou H, Zhao ZK. 2012. A multi-omic map of the lipid-producing yeast *Rhodosporidium toruloides*. Nat Commun 3:1112. doi:10.1038/ncomms2112.

45. Larsson C, Påhlman I-L, Ansell R, Rigoulet M, Adler L, Gustafsson L. 1998. The importance of the glycerol 3-phosphate shuttle during aerobic growth of *Saccharomyces cerevisiae*. Yeast 14:347–357. doi:10.1002/(SICI)1097-0061(19980315)14:4<347:AID-YEA226>3.0.CO;2-9.

46. Vincent O, Carlson M. 1999. Gal83 mediates the interaction of the Snf1 kinase complex with the transcription activator Sip4. EMBO J 18:6672–6681. doi:10.1093/emboj/18.23.6672.

47. Roth S, Kumme J, Schüller H-J. 2004. Transcriptional activators Cat8 and Sip4 discriminate between sequence variants of the carbon source-responsive promoter element in the yeast *Saccharomyces cerevisiae*. Current Genetics 45:121–128. doi:10.1007/s00294-003-0476-2.

48. Galazka JM, Tian C, Beeson WT, Martinez B, Glass NL, Cate JHD. 2010. Cellodextrin transport in yeast for improved biofuel production. Science 330:84–86. doi:10.1126/science.1192838.

49. Pertea M, Kim D, Pertea GM, Leek JT, Salzberg SL. 2016. Transcript-level expression analysis of RNA-seq experiments with HISAT, StringTie and Ballgown. Nat Protoc 11:1650–1667. doi:10.1038/nprot.2016.095.

50. Block H, Maertens B, Spriestersbach A, Brinker N, Kubicek J, Fabis R, Labahn J, Schäfer F. 2009. Chapter 27 Immobilized-Metal Affinity Chromatography (IMAC), p. 439–473. *In* Burgess R (ed), Guide to protein purification, 2^nd^ ed., vol. 463. Elsevier, Amsterdam.

51. Buchanan CL, Connaris H, Danson MJ, Reeve CD, Hough DW. 1999. An extremely thermostable aldolase from *Sulfolobus solfataricus* with specificity for non-phosphorylated substrates. Biochem J 343 Pt 3:563–570.

52. Sievers F, Wilm A, Dineen D, Gibson TJ, Karplus K, Li W, Lopez R, McWilliam H, Remmert M, Söding J, Thompson JD, Higgins DG. 2011. Fast, scalable generation of high-quality protein multiple sequence alignments using Clustal Omega. Mol Syst Biol 7:539. doi:10.1038/msb.2011.75.

53. Li W, Cowley A, Uludag M, Gur T, McWilliam H, Squizzato S, Park YM, Buso N, Lopez R. 2015. The EMBL-EBI bioinformatics web and programmatic tools framework. Nucleic Acids Res 43:W580–4. doi:10.1093/nar/gkv279.

54. McWilliam H, Li W, Uludag M, Squizzato S, Park YM, Buso N, Cowley AP, Lopez R. 2013. Analysis Tool Web Services from the EMBL-EBI. Nucleic Acids Res 41:W597–600. doi:10.1093/nar/gkt376.

55. Saier MH, Reddy VS, Tsu BV, Ahmed MS, Li C, Moreno-Hagelsieb G. 2016. The Transporter Classification Database (TCDB): recent advances. Nucleic Acids Res 44:D372–9. doi:10.1093/nar/gkv1103.

56. Saier MH, Reddy VS, Tamang DG, Västermark A. 2014. The transporter classification database. Nucleic Acids Res 42:D251–8. doi:10.1093/nar/gkt1097.

57. Saier MH, Yen MR, Noto K, Tamang DG, Elkan C. 2009. The Transporter Classification Database: recent advances. Nucleic Acids Res 37:D274–8. doi:10.1093/nar/gkn862.

58. Saier MH, Tran CV, Barabote RD. 2006. TCDB: the Transporter Classification Database for membrane transport protein analyses and information. Nucleic Acids Res 34:D181–6. doi:10.1093/nar/gkj001.

59. Finn RD, Coggill P, Eberhardt RY, Eddy SR, Mistry J, Mitchell AL, Potter SC, Punta M, Qureshi M, Sangrador-Vegas A, Salazar GA, Tate J, Bateman A. 2016. The Pfam protein families database: towards a more sustainable future. Nucleic Acids Res 44:D279–85. doi:10.1093/nar/gkv1344.

60. Kumar A, Henrissat B, Arvas M, Syed MF, Thieme N, Benz JP, Sørensen JL, Record E, Pöggeler S, Kempken F. 2015. De novo assembly and genome analyses of the marine-derived *Scopulariopsis brevicaulis* strain LF580 unravels life-style traits and anticancerous scopularide biosynthetic gene cluster. PLoS ONE 10:e0140398. doi:10.1371/journal.pone.0140398.

61. Mojzita D, Penttilä M, Richard P. 2010. Identification of an L-arabinose reductase gene in Aspergillus niger and its role in L-arabinose catabolism. J Biol Chem 285:23622–23628. Doi:10.1074/jbc.M110.113399.

